# Epigenetic feedback and stochastic partitioning during cell division can drive resistance to EMT

**DOI:** 10.1101/2020.04.29.068239

**Authors:** Wen Jia, Shubham Tripathi, Priyanka Chakraborty, Adithya Chedere, Annapoorni Rangarajan, Herbert Levine, Mohit Kumar Jolly

## Abstract

Epithelial-mesenchymal transition (EMT) and its reverse process mesenchymal-epithelial transition (MET) are central to metastatic aggressiveness and therapy resistance in solid tumors. While molecular determinants of both processes have been extensively characterized, the heterogeneity in the response of tumor cells to EMT and MET inducers has come into focus recently, and has been implicated in the failure of anti-cancer therapies. Recent experimental studies have shown that some cells can undergo an irreversible EMT depending on the duration of exposure to EMT-inducing signals. While the irreversibility of MET, or equivalently, resistance to EMT, has not been studied in as much detail, evidence supporting such behavior is slowly emerging. Here, we identify two possible mechanisms that can underlie resistance of cells to undergo EMT: epigenetic feedback in ZEB1/GRHL2 feedback loop and stochastic partitioning of biomolecules during cell division. Identifying the ZEB1/GRHL2 axis as a key determinant of epithelial-mesenchymal plasticity across many cancer types, we use mechanistic mathematical models to show how GRHL2 can be involved in both the abovementioned processes, thus driving an irreversible MET. Our study highlights how an isogenic population may contain subpopulation with varying degrees of susceptibility or resistance to EMT, and proposes a next set of questions for detailed experimental studies characterizing the irreversibility of MET/resistance to EMT.

## Introduction

Epithelial-Mesenchymal Transition (EMT) is a cell biological process involved in driving cancer metastasis and therapy resistance – the two grand clinically unsolved challenges. EMT and its reverse process Mesenchymal-Epithelial Transition (MET) are believed to enable cancer cell dissemination from the primary tumor, facilitate survival in the bloodstream, and are implicated in extravasation and the formation of macrometastases at multiple distant organs [1]. Thus, understanding the dynamics of EMT and MET is essential to develop novel therapeutic interventions.

Recent studies have highlighted that EMT and MET are not binary processes as thought earlier [1]. Instead, besides the epithelial and mesenchymal phenotypes, cells can acquire and stably maintain one or more hybrid epithelial/mesenchymal (E/M) phenotypes. These hybrid E/M phenotypes may drive collective cell migration as clusters of tumor cells (CTCs) and can be more aggressive than cells in pure epithelial or mesenchymal phenotypes [2]. Importantly, tumor cells may switch among different phenotypes - E, M and hybrid E/M [3–5]. Such dynamic and reversible switching can help tumor cells to overcome various challenges during disease progression such as anoikis [6], and assaults by the immune system [7]. Thus, epithelial-mesenchymal plasticity (EMP) – a combination of EMT and MET – needs to be utilized with spatiotemporal precision to drive metastasis. For example, a failure to undergo MET at a metastatic site may compromise colonization [8,9]. The reversibility of EMT and MET, mediated by multiple interconnected feedback loops regulating a balance between epithelial and mesenchymal traits, thus forms the backbone of metastasis [10].

Are EMT and MET always reversible? Recent experiments decoding the dynamics of EMT/MET using live-cell imaging and/or induction and withdrawal of various EMT-inducing external signals such as TGFβ or tuning the levels of EMT-specific transcription factors (EMT-TFs) have provided important insights into the reversibility of EMT and MET. Cells induced to undergo EMT for shorter durations (~2-6 days) may revert to an epithelial state after withdrawal of the signal/stimulus. However, some cells exposed to EMT-inducing signals for longer durations (~10 days or more) may get ‘locked’ in a mesenchymal state, making EMT largely irreversible, at least for the timescale observed experimentally [11–14]. The possibility of an irreversible EMT is also supported by multiple phenomenological observations [15–17]. Multiple mechanisms have been proposed to explain the existence of a ‘tipping point’ – a time point beyond which continued treatment with EMT inducing signals can drive an irreversible EMT. These include self-stabilizing feedback loops [18–21] in the regulatory circuits for EMT/MET and/or epigenetic alterations [13,22,23]. However, similar investigations about the irreversibility of MET, or in other words, the resistance of epithelial cells to undergo EMT in response to EMT-inducing signals, remain to be done. Some sporadic observations about the resistance of epithelial cells to undergo EMT have been reported [14,24], but a causative mechanistic understanding still remains elusive.

Here, we propose two independent mechanism that may explain the resistance of epithelial tumor cells to undergo EMT: first, epigenetic feedback mediated via GRHL2 – an MET-inducing factor [25–27], and second, stochastic partitioning of parent cell biomolecules among the daughter cells at the time of cell division [28–30]. GRHL2 and miR-200 both form mutually inhibitory feedback loops with ZEB1 – a key EMT-TF [31]. Previously, we have shown that incorporating an epigenetic feedback term acting on the inhibition of miR-200 by ZEB1 could drive an irreversible EMT [13]. This epigenetic feedback term was incorporated at a phenomenological level to represent the idea that the longer a gene is turned on, the easier it becomes for it to stay transcriptionally active; thus, epigenetic feedback modulated the thresholds for the influence of a transcription factor on its downstream target [32,33]. Conversely, here, we show that incorporating this epigenetic feedback loop acting on the inhibition of ZEB1 by GRHL2 can cause an irreversible MET. Cells undergoing irreversible MET may exhibit resistance in undergoing EMT when exposed to EMT-inducing signal. Thus, our results offer a conceptual framework to decode the impact of various epigenetic mechanisms in terms of modulating the reversibility of EMT and MET in a cancer cell population. Further, our previous analysis illustrated how epithelial-mesenchymal heterogeneity can be generated from a phenotypically homogeneous population by stochastic partitioning of molecules during cell division [3]. Here, we demonstrate how this stochasticity in the presence of GRHL2 can lead to an irreversible MET or, in other words, a resistance to undergo EMT. Together, our results describe how an isogenic cellular population may contain these different subpopulations with varying degrees of susceptibility and resistance to undergo EMT in response to an EMT-inducing signal.

## Results

### GRHL2 correlates with a more epithelial phenotype across many cancer types

GRHL2 has been identified as an MET inducer in breast cancer [25–27], where it forms a mutually inhibitory feedback loop with ZEB1, an EMT-TF. Overexpression of GRHL2 suppresses EMT induced by TGF-β or Twist by directly binding to the ZEB1 promoter, and inhibits various other properties associated with a partial or complete EMT such as higher mammosphere-forming efficiency and anoikis resistance [6,25,34,35]. In general, the GRHL2/ZEB1 feedback loop was identified as a key regulator of EMP and associated traits in breast cancer, lung cancer [36], colorectal cancer [37] and ovarian cancer [38,39]. Consistently, GRHL2 was shown to inhibit EMT in gastric cancer [40], oral cancer [41] and pancreatic cancer [42]. These reports drove us to investigate the correlation of levels of GRHL2 with EMT/MET across many cancer types both in the Cancer Cell Line Encyclopedia (CCLE) cohort [43] and in many TCGA datasets.

GRHL2 levels correlated positively with CDH1 (E-cadherin) levels and negatively with ZEB1 in the CCLE dataset and TCGA datasets from breast cancer, ovarian cancer and colorectal cancer (Fig 1). Given that GRHL2 is one of the top transcriptional activators of CDH1 and ZEB1 is one of its strongest transcriptional repressors, ZEB1 and CDH1 correlated negatively (Fig S1). We also investigated the correlation of GRHL2 levels with three transcriptomics-based EMT scoring algorithms – MLR, KS, and 76GS. While MLR and KS methods assign higher scores to mesenchymal samples, the 76GS method assigns higher scores to epithelial samples [44]. As expected, GRHL2 levels correlated positively with EMT scores calculated via the 76GS method and negatively with EMT scores calculated by MLR or KS methods across these TCGA datasets (Fig S1). Consistently, a constitutive expression of GRHL2 in an inducible EMT model system – HMLE cells that contain a Twist-ER fusion – led to reduced ZEB1 levels and corresponding changes in EMT scores as identified by all three abovementioned EMT scoring metrics (Fig S2).

**Figure 1:**
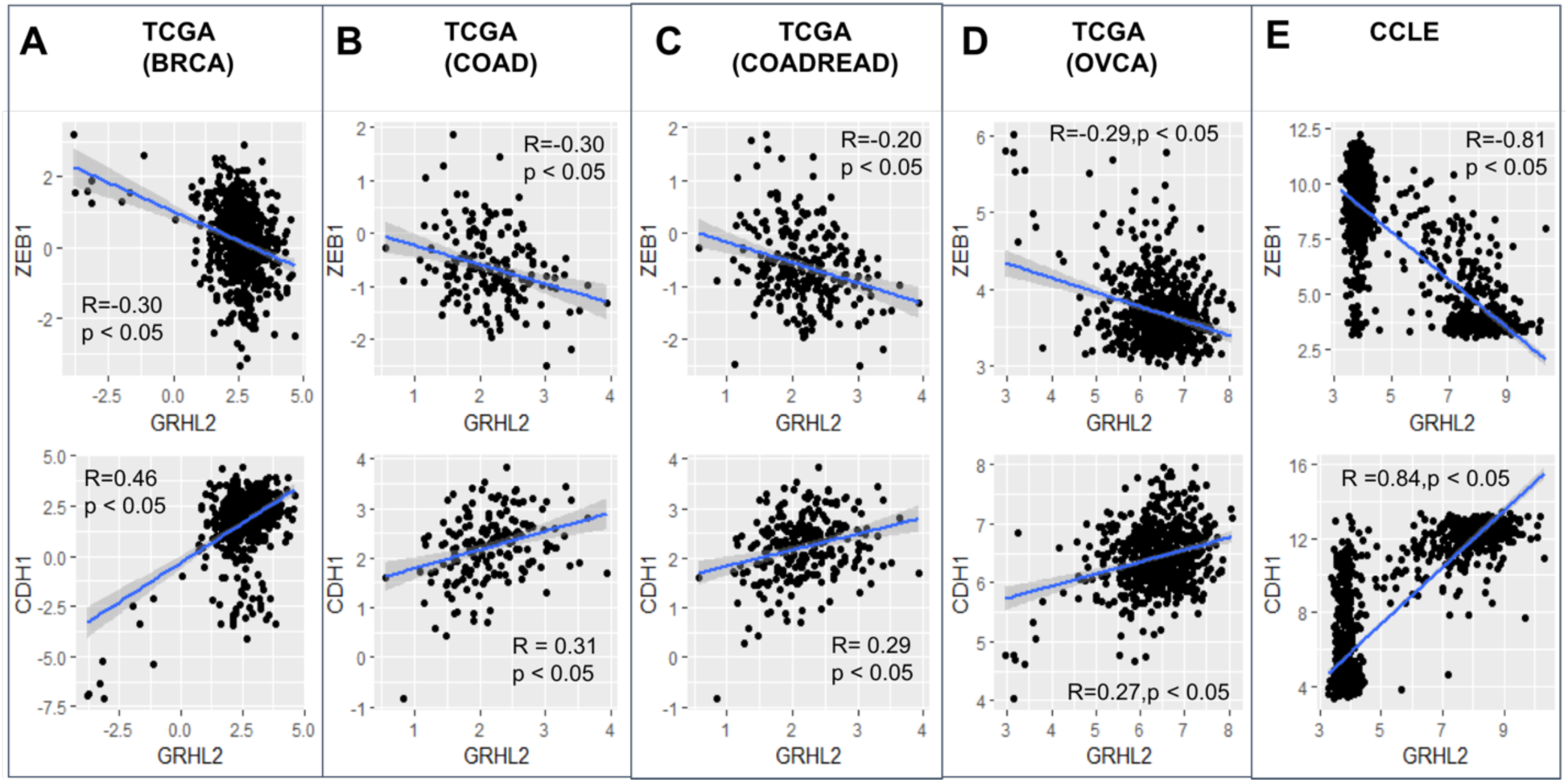
GRHL2 correlates with an epithelial phenotype. Scatter plots showing correlation of GRHL2 with EMT-TF ZEB1 and CDH1 (E-cadherin) in TCGA datasets and CCLE – A) breast cancer, B) colon adenocarcinoma, C) colorectal adenocarcinoma, D) ovarian carcinoma, E) CCLE. R, p denote Pearson’s correlation coefficient and corresponding p-value for corresponding plot.

Next, in the CCLE dataset, we calculated the pairwise correlations of various canonical epithelial and mesenchymal markers and regulators. We found that GRHL2 correlates positively with its family members GRHL1 and GRHL3, its downstream target OVOL2 and corresponding family member OVOL1, and with CDH1, while it correlates negatively with TWIST1/2, SNAI1, VIM, and ZEB1/2 (Fig 2A, S3A). On the other hand, ZEB1 correlates negatively with GRHL1/2/3, OVOL1/2 and positively with SNAI1/2, TWIST1/2 and VIM (Fig 2A). With these consistent observations regarding the antagonistic roles of ZEB1 and GRHL2 in regulating EMT/MET, we next calculated the correlation of all genes in CCLE with GRHL2 and with ZEB1, and observed that most genes that showed significant correlation with both of them were either positively correlated with GRHL2 and negatively with ZEB1 or *vice-versa*. The relatively smaller set of genes that correlated either positively or negatively with both ZEB1 and GRHL2 showed relatively weak correlations (r <0.3) (Fig 2B, S3B). Together, these observations indicate that GRHL2 associates with epithelial traits across cancer types.

**Figure 2:**
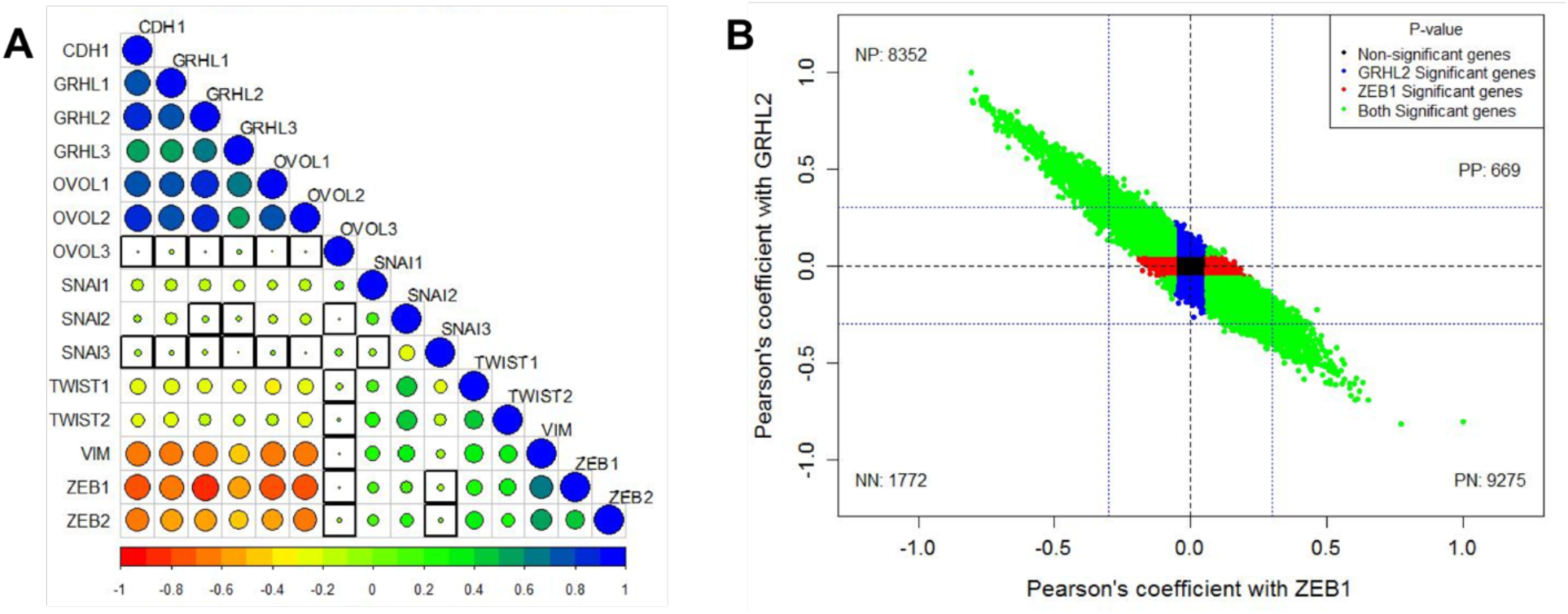
GRHL2/ZEB1 axis correlates with EMT/MET across cancer types. A) Pairwise Pearson’s correlation between different EMT and MET regulatory genes in the CCLE dataset. Pearson’s correlation value (cor) of each gene pair is represented as the size of the circle and filled with corresponding color from the color palette represented below the ranging from −1(red) to +1(blue). Boxes highlighted by the black squares represent insignificant (p>0.01) correlation. B) Scatter plot of genes correlated using Pearson correlation method with GRHL2 and ZEB1 in CCLE dataset. Each dot represents one gene and coordinates are Pearson cor values with ZEB1 and with GRHL2. Color of the dots is based on the p-value obtained from correlation test, Blue dots for genes having p<0.05 with GRHL2 and p>0.05 with ZEB1. Red dots for genes having p<0.05 with ZEB1 and p>0.05 with GRHL2. Green dots for genes having p<0.05 with GRHL2 and ZEB1. Black dots for genes having p>0.05 05 with GRHL2 and ZEB1. Numbers in each quadrant represent the number of genes in that quadrant.

### Epigenetic feedback on self-activation of GRHL2 does not largely affect EMT/MET dynamics

We have previously analyzed the dynamics of the EMT/MET regulatory network that incorporates the connection of GRHL2 with the two double negative loops that are central to EMT/MET dynamics: miR-200/ZEB1 and miR-34/SNAIL [36] (Fig 3A) miR-34 and miR-200 are EMT-inhibiting microRNAs that can inhibit the translation of EMT-TFs SNAIL and ZEB1, thus safeguarding an epithelial phenotype. ZEB and SNAIL can repress E-cadherin and other epithelial genes, and/or drive the expression of mesenchymal genes [31,45]. The knockdown of GRHL2 drives EMT and impairs collective cell migration, while its overexpression may drive an MET [46], the reverse is true for ZEB1 [31,47]. Finally, ZEB1 and GRHL2 can both promote their own expression, albeit indirectly [31,48].

**Figure 3:**
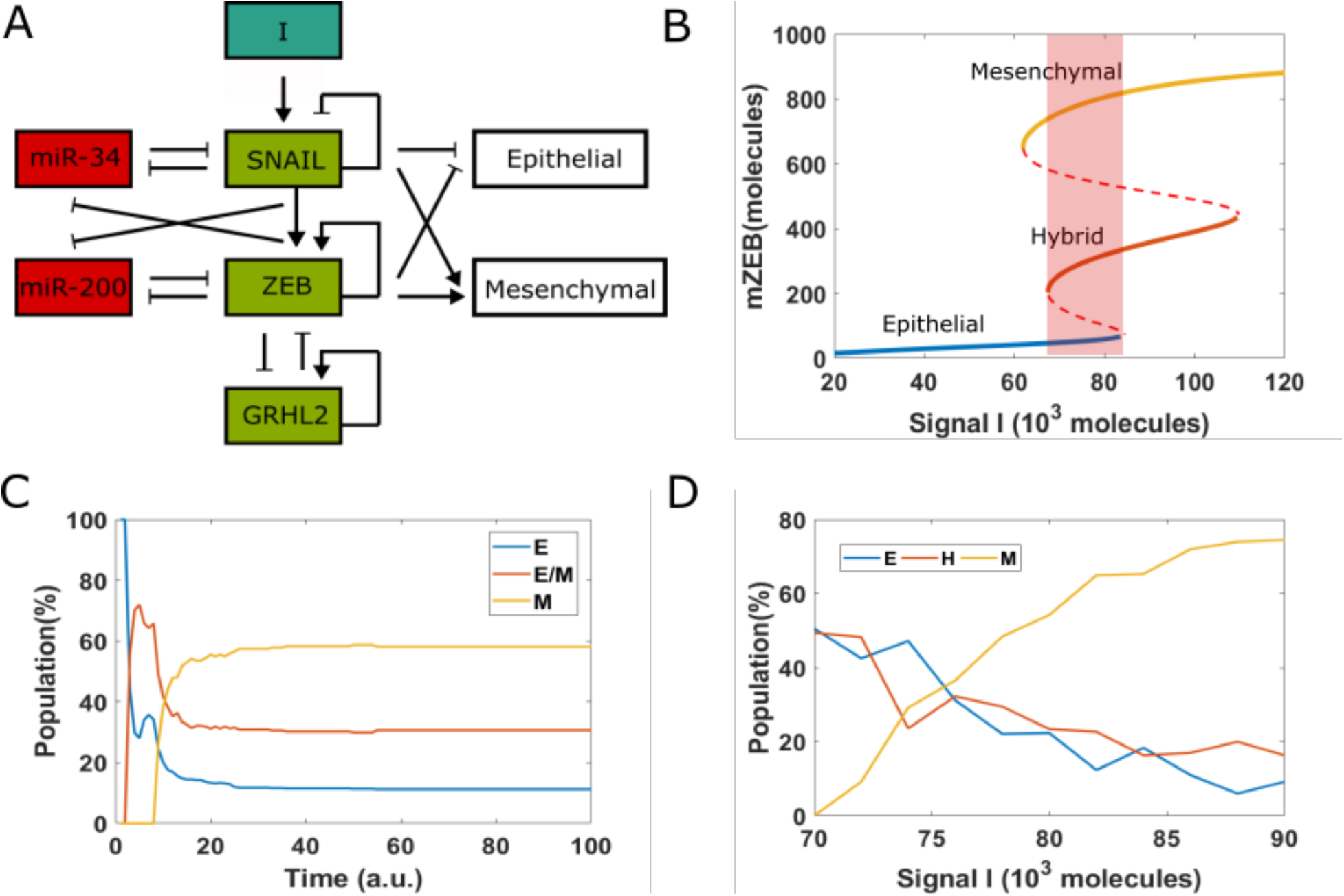
EMT decision-making network. A) A core network regulating EMT via two mutually inhibition loops between miR-34 (miR-200) and SNAIL (ZEB). Signal I represents external EMT-inducing signals such as HGF, TGF-β, NF-κB and HIF1α, among others. GRHL2 is a transcription factor which forms a mutually inhibitory loop with ZEB. (B) Bifurcation diagram (shown for the levels of ZEB mRNA) with I as the bifurcation parameter. Solid lines represent stable states, i.e. epithelial, hybrid or mesenchymal state, and dashed lines represent unstable states. Shaded rectangle represents the values of I for which all three phenotypes can co-exist. C) Starting from epithelial state (miR-200=17,000, mZEB=50, ZEB=10,000 molecules) with different pre-fixed threshold value of inhibition of ZEB by GRHL2, 100 cells are treated by fixed high EMT-inducing signal (I=75,000 molecules). D) Population distribution changes as a function of signal I.

The bifurcation diagram (Fig 3B) shows that this network can enable cells to exhibit three distinct phenotypes: epithelial (E), hybrid epithelial/mesenchymal (H) and mesenchymal (M). At low levels of an external EMT-inducing signal I, cells are in an epithelial phenotype; they switch to a hybrid E/M phenotype and finally a mesenchymal phenotype as the levels of I increase. For certain range of values of I, more than one phenotype may co-exist, enabling stochastic cell-fate transitions. Under these conditions, the phenotype exhibited by a cell may switch spontaneously under the influence of biological noise (Fig 3B) [49], leading to non-trivial phenotypic distributions.

Recent experiments showed how cells in an isogenic population can exhibit varying degrees of susceptibility or resistance to EMT in response to EMT induction [14]. A large subset of single-cell clones (SCCs) (72%) derived from this population successfully underwent EMT and displayed irreversible EMT upon withdrawal of the signal. The remaining 28% SCCs, in response to EMT inducing signal, did not exhibit a reduction in epithelial traits and only underwent a reversible and partial EMT. To capture this behavior in our simulations, we started from all cells in an epithelial state, but with each of them with different random values of threshold of the Hills function denoting the transcriptional inhibition of ZEB by GRHL2. Thus, in this simulation, each cell in this population has different extents of inhibition of ZEB by GRHL2. For a fixed value of EMT-inducing signal I = 75*10^3^, in 100 heterogenous cells, only 58% of them underwent an EMT (Fig 3C). This percentage was higher for larger values of I, but a small subset of cells remained in H or E state, confirming that some cells can intrinsically resist EMT-inducing signals (Fig 3D).

To depict these transitions, we plotted the stochastic dynamics of a population of 1000 cells. The mean value of the signal was fixed at 75*10^3^ molecules and all cells were initially in an epithelial state. We observed a stable phenotypic distribution with 58% E, 37% hybrid E/M and 5% M cells (Fig 4B). Next, we added a strong epigenetic feedback governing the threshold value of GRHL2 that governs its self-activation. The threshold value, instead of being a static variable, is now governed by a dynamical equation with a lower steady-state value which is proportional to the feedback factor α. This feedback has a minimal effect on the bifurcation diagram of the circuit (compare the black and blue curves in Fig 4A); this feature is true for all biologically relevant values of α. We next investigated the effect of this feedback on the equilibrium phenotypic distribution. We started with a scenario where the entire population (n=1200) exhibits an epithelial phenotype, and tracked the dynamics when a noise term was added to induce spontaneous transitions among the different states (SI sections 1-3). We notice that the phenotypic distribution seen for the case with epigenetic feedback (55% E, 37% H and 8% M) is largely similar to the scenario without any such feedback (58% E, 37% H and 5% M) (Fig 4B).

**Figure 4:**
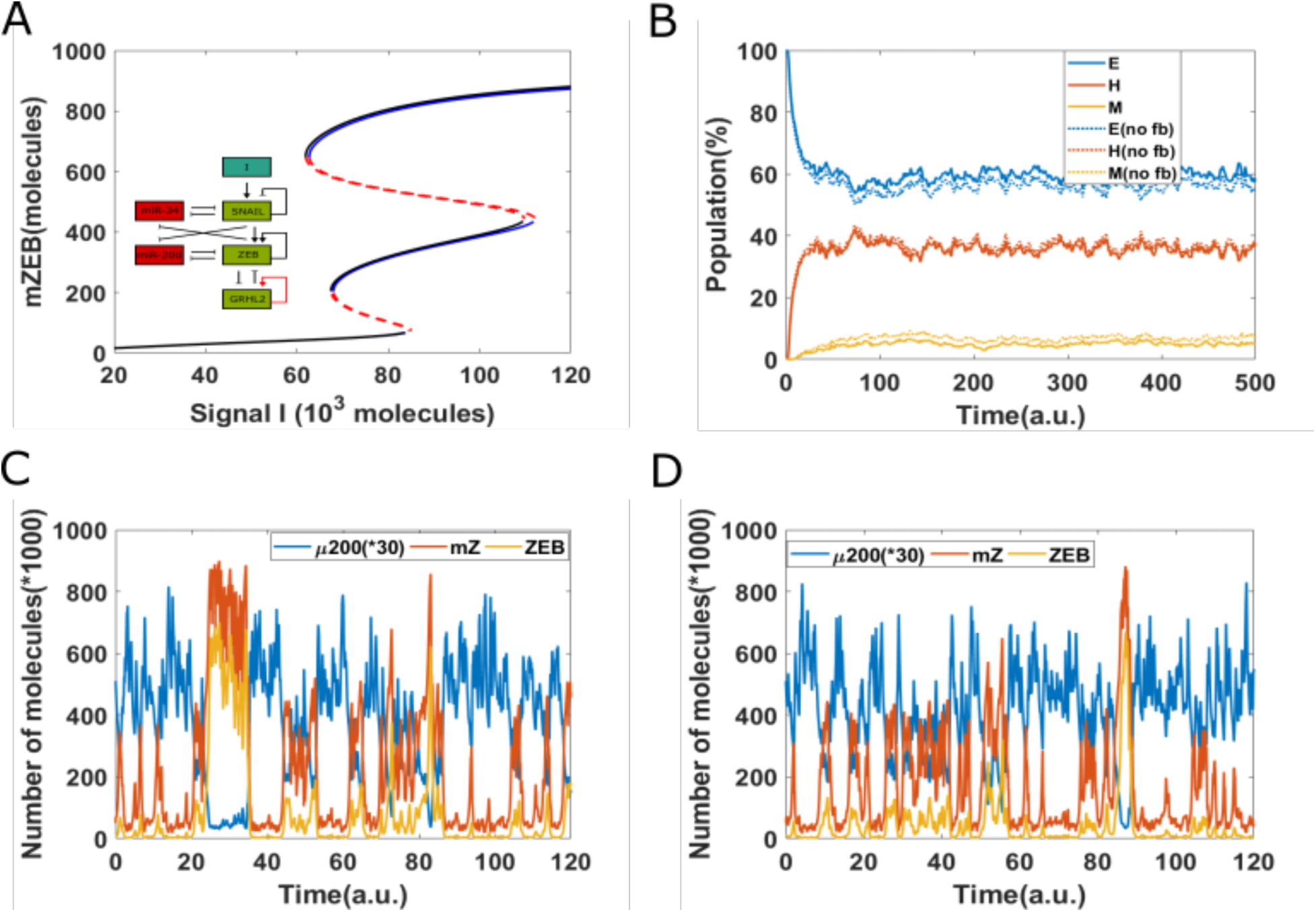
Epigenetic feedback on GRHL2 self-activation. A) The bifurcation diagrams for core EMT circuit with/without epigenetic feedback on self-activation of GRHL2. Black curve denotes the case without any epigenetic feedback; blue curve represents the epigenetic feedback case. B) Starting from 100% cells in an epithelial state (miR-200=17,000, mZEB=50, ZEB=10,000 molecules), simulation results showing how the population changes as a function of simulation time, on addition of noise. Dashed lines represent no epigenetic feedback case, solid lines represent case with strong epigenetic feedback (a = 0.22) on GRHL2’s self-activation (Signal I_0_=75,000 molecules). (C) A representative dynamical trajectory for no epigenetic feedback case. (D) A representative dynamical trajectory for strong epigenetic feedback case.

These results strengthen the observations made from the bifurcation diagram. Moreover, a comparative analysis of the dynamics of two cases show minimal differences – in both cases, instances of partial EMT/MET and complete EMT/MET can be observed, and cells can continue to transition among all three phenotypes (Fig 4C,D). Together, these results suggest that epigenetic feedback acting only on self-activation of GRHL2 has only a weak effect on the dynamics of EMT/MET.

### Epigenetic feedback on the inhibition of ZEB by GRHL2 can stabilize an epithelial state

Next, we examined the effect of adding an epigenetic feedback on the inhibition of ZEB by GRHL2. Unlike the scenario of epigenetic feedback on GRHL2 self-activation, incorporating epigenetic feedback on the inhibition of ZEB by GRHL2 significantly alters the bifurcation diagram.

Compared to the case without any epigenetic feedback (black curve), the bifurcation curve for a case of strong epigenetic feedback case (blue lines) shifts to the right (Fig 5A), suggesting that a higher external stimuli is required to induce EMT. In other words, this epigenetic feedback can stabilize an epithelial state, or in other words, offer resistance to undergo an EMT. Consistently, large changes were noted in the population distribution as well - the equilibrium population distribution for the case including epigenetic feedback on inhibition of ZEB by GRHL2 was 79% E, 20% H and 1%M; thus, compared to the control case, the epithelial population increased by 22%, while H and M populations both decreased (Fig 5B). Moreover, corresponding dynamical trajectories show that in the presence of a strong epigenetic feedback on inhibition of ZEB by GRHL2, it becomes exceedingly difficult for cells to reach a mesenchymal state. Thus, most cells stay much more robustly in an epithelial state (Fig 5C-D). Hence, a sufficiently strong epigenetic feedback on the inhibition of ZEB by GRHL2 can make cells resistant to undergoing a full EMT.

**Figure 5:**
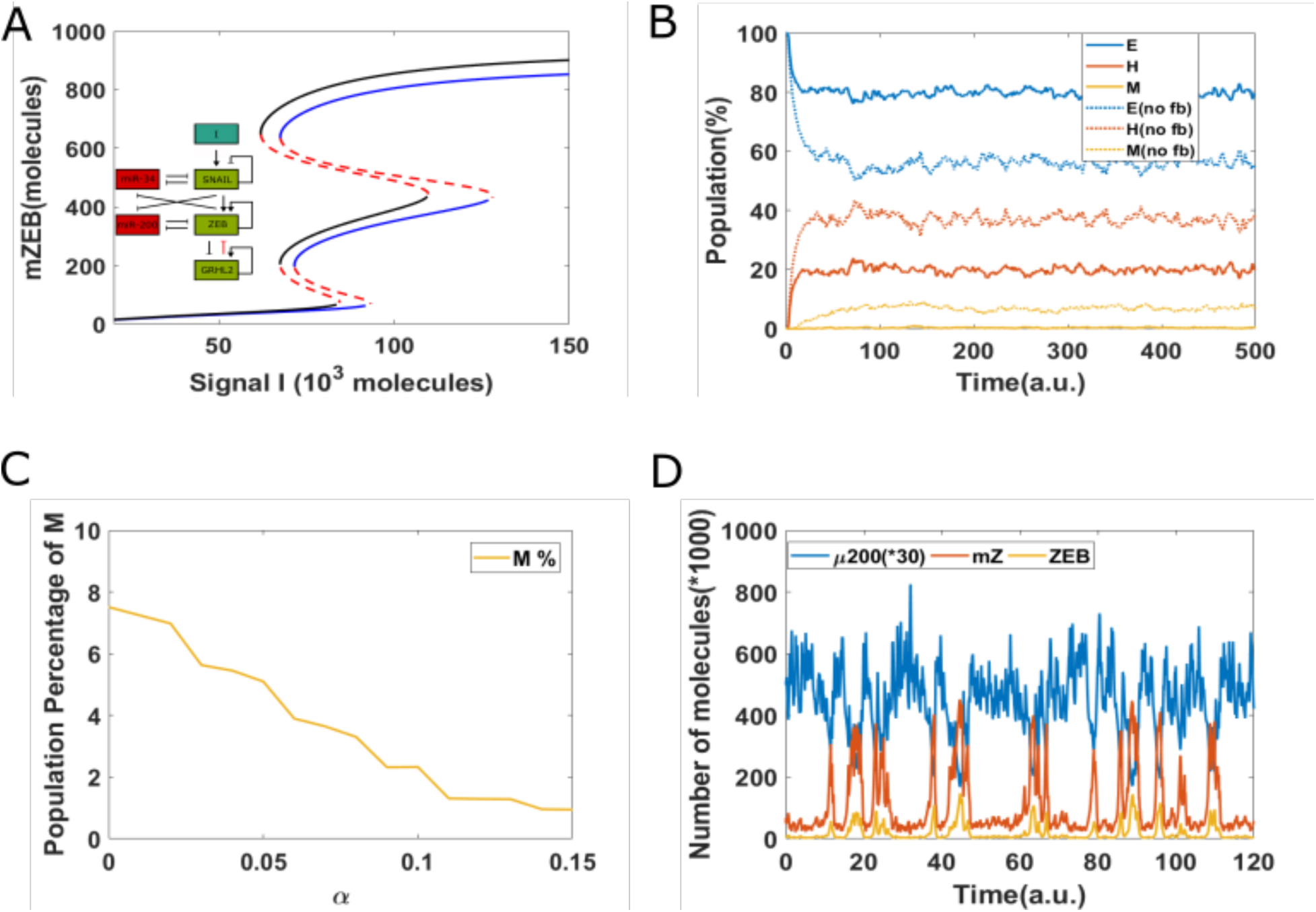
Epigenetic feedback on inhibition of ZEB by GRHL2. A) The bifurcation diagrams for core EMT circuit with/without epigenetic feedback on the inhibition of ZEB by GRHL2. Black curve represents the case without any epigenetic feedback; blue curve represents the epigenetic case. B) Starting from all cells in an epithelial state (miR-200=17,000, mZEB=50, ZEB=10,000 molecules), simulation results showing how the population changes as a function of simulation time. Dashed lines represent no epigenetic feedback case, and solid lines represent case with strong epigenetic feedback (α = 0.14) on the inhibition of ZEB by GRHL2 (Signal I_0_=75,000 molecules). (C) The percentage of population which exhibit M phenotype, for varying values of α. (D) A sample dynamical diagram for strong feedback case.

The abovementioned analysis was also conducted on the EMT circuit without miR-34 (because miR-34/ SNAIL has been proposed to be a noise-buffering integrator for EMT). In this simplified model, SNAIL serves as inducing signal. The simulation showed similar results that epigenetic feedback of GRHL2 on the inhibition of ZEB can stabilize an epithelial state, while the epigenetic feedback on GRHL2’s self-activation does not largely change EMT/MET dynamics (SI section 5; Fig S4-S6).

Next, we studied whether adding an epigenetic feedback on the inhibition of ZEB by GRHL2 can affect the reversibility of EMT. To mimic the experiment where cells are treated with an EMT-inducing signal (say TGF-β; represented by I in simulations) for varying time durations, here, we increased the value of I = 125*10^3^ molecules and then decreased it back to original value of I =71*10^3^ molecules. In the case without any epigenetic feedback case, for a short duration increase in the levels of I, cell can quickly revert to epithelial state upon the removal of the signal (Fig. 6A). For somewhat longer treatment, the cell can stay in hybrid E/M phenotype for a long time and eventually revert to being epithelial (Fig. 6B). A further increase in the duration of cells to EMT-induction can render the induced EMT irreversible; i.e. cells can stay in mesenchymal state and not revert to being epithelial even after the EMT-inducing signal is effectively withdrawn. However, the presence of epigenetic feedback on the inhibition of ZEB by GRHL2 can alter the abovementioned dynamics. In presence of such feedback, a cell can revert to being epithelial rapidly soon after the external signal is withdrawn, irrespective of the duration for which cells were exposed to this signal (Fig 6D-F). All these results reveal that epigenetic feedback on the inhibition of ZEB by GRHL2 can significantly stabilize an epithelial state and thus increase the reversibility of EMT.

**Figure 6.**
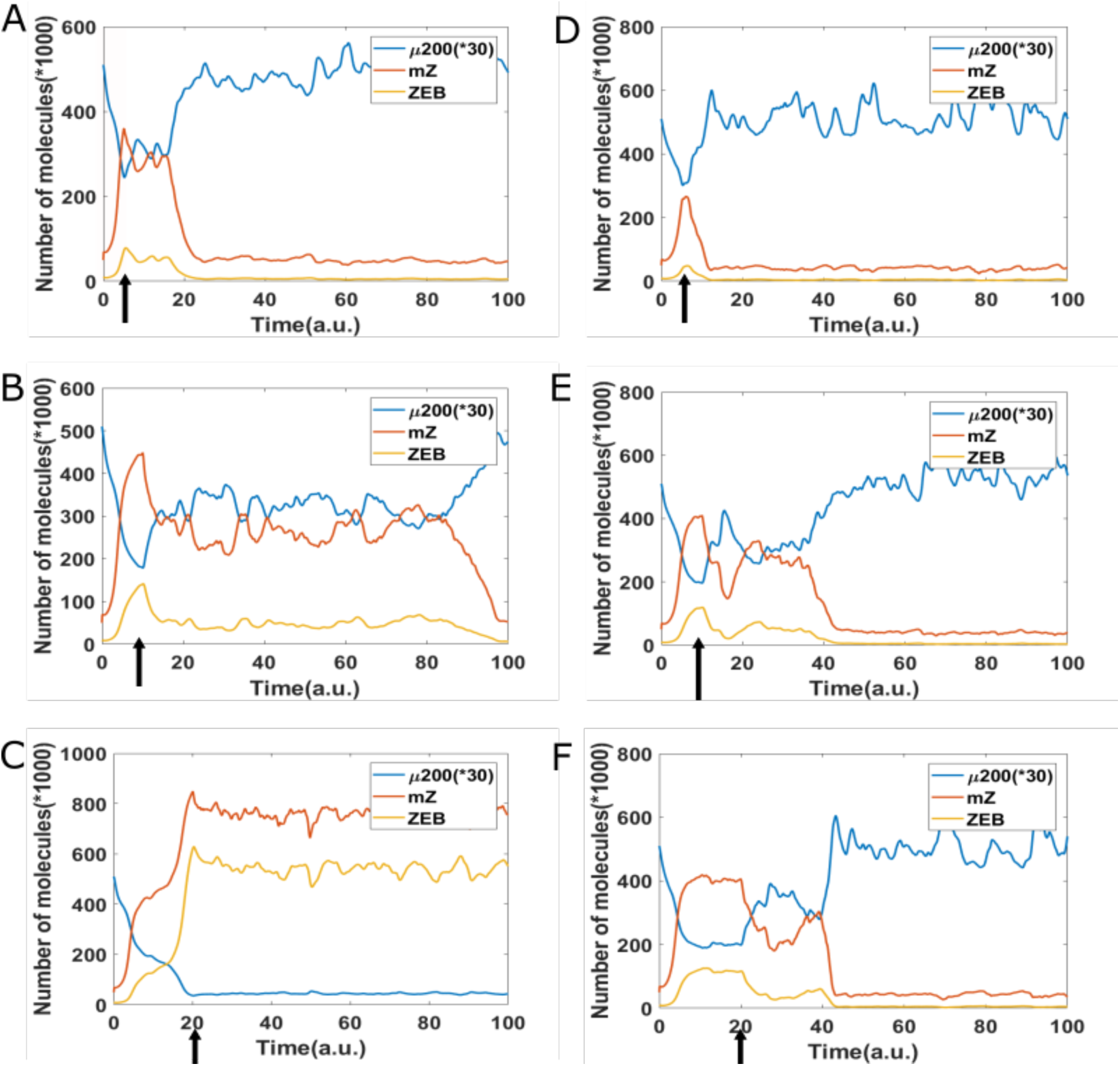
Reversibility of EMT. Starting from epithelial state (miR-200=17,000, mZEB=50, ZEB=10,000 molecules), a cell is treated by different time duration (5, 10, 20 arbitrary units (a.u.), as marked by arrow) of high EMT-inducing signal (I=125,000 molecules), corresponding to the {H, M} bistable region. Then, this signal is reduced to a lower level (I=71,000 molecules) corresponding to the {E, M} bistable region. Panels A-C represents the case without epigenetic feedback, and D-F represents the case with strong epigenetic feedback on inhibition of ZEB by GRHL2.

### Noise in the partitioning of parent cell biomolecules among the daughter cells can cause a seemingly irreversible MET at the population level

So far, we have described an epigenetic-based mechanisms which may underlie irreversibility of MET. We next investigated the effect of another stochastic behavior in population of cancer cells – partitioning of molecules during cell division [28–30] – independent of epigenetic feedback. Thus, we investigated the dynamics of EMT/MET at a population level, where we incorporated cell division with an average doubling time of 38 hours, typical of cancer cells [50]. At every instance of cell division, we incorporated some noise in the partitioning of parent cell biomolecules among the daughter cells.

Our previous analysis has shown that such noise in the distribution of biomolecules during cancer cell division can generate epithelial-mesenchymal heterogeneity in an initially homogeneous population of cancer cells [3]. Thus, a daughter cell may or not have the same phenotype (E, hybrid E/M or M) as the parent cell. Further, this noisy partitioning can lead to some cells in the population undergoing a seemingly irreversible EMT if the cells are treated with an EMT-inducing signal followed by the withdrawal of the EMT-inducing signal. We investigated if the noisy partitioning model described previously can also lead to a seemingly irreversible MET at the population level.

Starting with a population of all mesenchymal cells on day 0 (all cells had high concentration of the EMT-inducing signal initially), fixed dosages of the EMT-inducing signal were withdrawn each day for a period of 10 days (to simulate MET). Day 11 onwards, fixed dosages of the EMT-inducing signal were added for another 10 days. We carried out the simulation in the absence and presence of GRHL2, and in both cases, ~40% of the cells had undergone MET by day 10. In the absence of GRHL2, upon treatment with the EMT-inducer starting on day 11, almost all the cells in the population that had undergone MET returned to a mesenchymal state by day 10. Thus, in this scenario, MET was reversible. However, in the scenario when GRHL2 was present, it was observed that >10% of cells still exhibited an epithelial phenotype on day 20 (Fig 7). These cells thus represented a sub-population that had under-gone an irreversible MET; i.e. a subpopulation which would exhibit resistance to EMT upon exposure to EMT-inducing signal.

**Figure 7.**
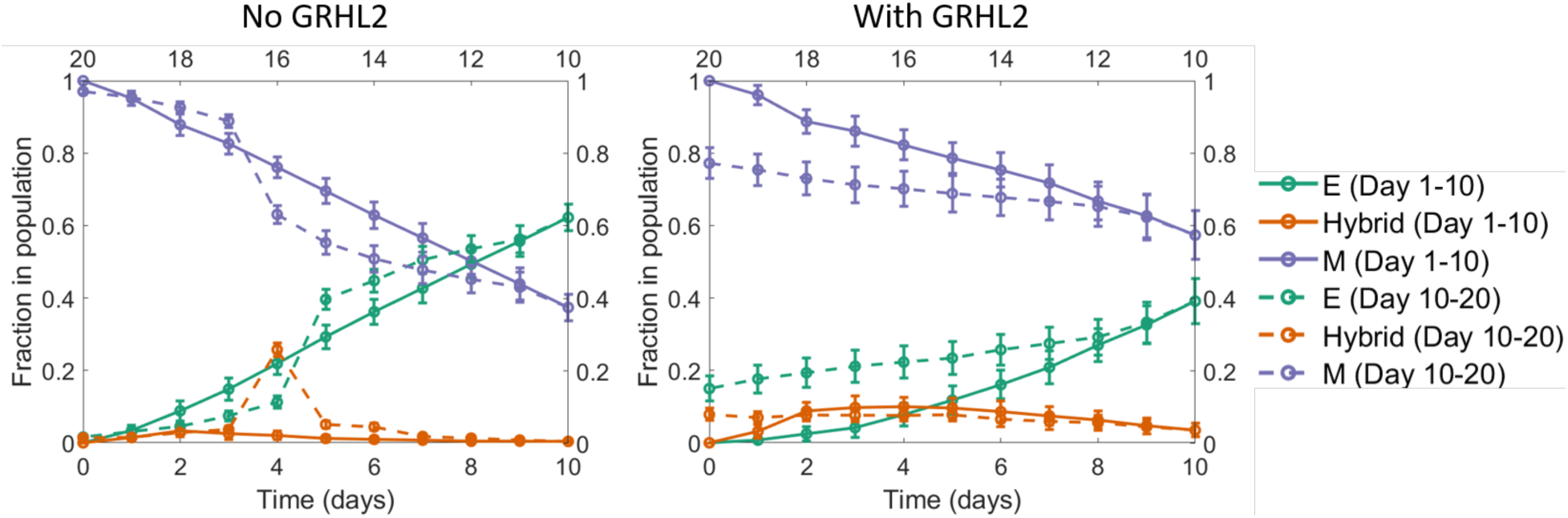
A seemingly irreversible MET at the population level. Starting with a population of mesenchymal cells on day 0 (all cells with very high value of the EMT inducer), the EMT inducer was withdrawn in fixed dosages each day for the first 10 days. As a result, a large fraction of cells in the population underwent MET. Day 11 onwards, fixed dosages of the EMT inducer were added each day for the next 10 days. In the absence of GRHL2 (left panel), the fraction of mesenchymal cells went back to nearly 100%, same as the value on day 0. However, in the presence of GRHL2 (right panel), ~15% of the cells in the population were epithelial on day 20. These cells thus represented a subpopulation that had undergone a MET that is irreversible at least on the time scale investigated here. The mean over 16 independent simulation runs is shown here. The error bars indicate the standard deviation over the independent runs.

## Discussion

With an increasing appreciation of the nonlinear dynamics of EMT/MET at the single-cell level [51–53] and its implications for metastatic aggressiveness [9], questions regarding the degree of reversibility/irreversibility of EMT/MET in different contexts and the molecular determinants of these processes have gained importance. Recent studies have illustrated that the trajectories taken by individual cells in the high-dimensional molecular and/or morphological landscape of EMP *en route* to EMT and MET may be different [51,54]; thus, MET cannot be simply thought of as a mirror image of EMT. There may be molecular and/or morphological changes happening at different stages of EMT/MET to varying degrees, hence making it difficult to identify the molecular mechanisms that may render the dynamics of EMT/MET as reversible or irreversible.

The dynamics of EMT has been studied much more in detail as compared to that of MET [51–56]; therefore, it is not surprising that irreversible EMT has been reported more frequently. The degree of reversibility of EMT has been proposed to be largely a function of the timescale of EMT induction and corresponding epigenetic changes. However, a causative role of epigenetic changes in regulating the irreversibility of EMT remains to be firmly established [11–14]. Here, we propose two independent mechanisms that may enable an irreversible MET – epigenetic feedback mediated by the inhibitory action of GRHL2 on the promoter of ZEB1, and noise in the partitioning of biomolecules during cell division.

GRHL2 can inhibit and reverse EMT and associated molecular and/or morphological traits. Specifically, overexpression of GRHL2 has been shown to induce epithelial gene expression, inhibit mesenchymal gene expression, restore metabolic reprogramming caused by EMT, and suppress tumor cell migration/invasion, at least in breast and ovarian cancer cell lines [46]. It can also activate directly or indirectly other drivers of an epithelial phenotype such as p63 [57,58], OVOL2 [59,60], and miR-200 [38,61]. While epigenetic changes induced by EMT-TFs have been reported extensively [62], recent studies have pointed out the possibility of GRHL2 contributing to epigenetic control of genes involved in EMT/MET [39]. GRHL2 can employ both DNA methylation and histone modification to inhibit and/or reverse EMT, and also act as a pioneer transcription factor that can regulate chromatin accessibility at epithelial enhancers [39,63,64].

The experimental observations described above offer a possible underlying mechanism by which GRHL2 overexpression can resist EMT, as demonstrated by our model simulations. Indeed, a global epigenetic program that limits the action of ZEB1 was found to underlie the retention of epithelial traits in cells exposed to persistent Twist1 activation for 21 days [14]. In single-cell clones established from an inducible HMLE-Twist1 population, two subsets responded quite differently (E-SCCs, M-SCCs); while both E-SCCs and M-SCCs exhibited an upregulation of mesenchymal markers to a similar degree upon Twist1 induction, but E-SCCs did not display any reduction in epithelial genes even after 28 days of Twist1 induction. Upon inactivation of Twist1, E-SCCs reverted to an epithelial phenotype, while M-SCCs did not. Changes in chromatin accessibility seen in E-SCCs also reverted upon Twist1 deactivation, but not in M-SCCs, suggesting a strong correlation of transcriptional changes induced in E-SCCs/M-SCCs and dynamic epigenetic status. Intriguingly, this study highlighted that isogenic/clonal cells can also exhibit variability in terms of their susceptibility or resistance to EMT/MET, indicating non-genetic mechanisms at play [65], as indeed captured via our simulations. Our simulations also offer a causal connection between the two axes phenomenologically associated with one another in this study – epigenetic changes driven by GRHL2 and consequent resistance to EMT.

More generally, resistance to EMT has also been observed across many cell lines, where their treatment with 5ng/ml TGFβ for 48 hours led to varying degrees of changes in molecular and morphological axes of EMT: spindle-shape acquisition, loss of junctional E-cadherin and/or ZO-1, and actin stress fiber formation [24]. Another recent study in multiple breast cancer cell lines showed that 100pM TGFβ treatment for 9 days need not be sufficient for at least a subset of cells within a cell line to lose their E-cadherin expression [52]. While longer-time measurements would probably be more appropriate to fully assess the degree of resistance to EMT (in other words, the degree of irreversibility of MET), both these studies emphasize the possible implications of non-genetic heterogeneity prevalent in multistable regulatory networks [66] as seen in EMT/MET and its associated traits such as stemness [67]. Different degrees of couplings between these multistable networks may enable co-occurrence of partial or full EMT with these associated traits. However, a mechanism-based mathematical modeling of these networks suggests that this association is not likely to be universal [68], hence offering a reconciliatory framework to integrate various contradictory results associating the epithelial, mesenchymal and hybrid E/M phenotypes with degrees of stemness [69–75].

Aside from epigenetic feedback, our results indicate one additional mechanism possibly contributing to irreversible MET (or resistance to EMT), namely stochasticity in the partitioning of molecules during cell division. Such noise in the distribution of molecules may affect cell-fate and drive non-genetic heterogeneity [28–30], leading to different phenotypic distributions in terms of EMT [3]. While EMT is believed to repress the cell cycle [76,77], this association remains controversial [78]. Thus, noise in the partitioning of parent cell biomolecules among the daughter cells can further alter the subpopulation structure, and may underlie different bimodal distributions of surface CDH1 expression seen in breast cancer cell lines [52].

Together, our results offer mechanistic insights into two possible mechanisms that may drive varying degrees of susceptibility and resistance to undergoing EMT in response to an EMT-inducing signal in a given isogenic population. Future efforts should decode the molecular mechanisms of any such epigenetic feedback of GRHL2 on ZEB1 expression as well as track the distribution of molecules during cell divisions happening while cells are being induced to undergo EMT/MET. Moreover, our study calls for concerted efforts to map the single-cell dynamics of MET induction.

## Acknowledgements

This work was supported by the National Science Foundation grants PHY-1427654 and PHY-1935762 (H.L.) and by the Ramanujan Fellowship (SB/S2/RJN-049/2018) awarded by the Science and Engineering Research Board, Department of Science and Technology, Government of India (M.K.J.).

## Materials and Methods

The analysis shown in Fig 1 and Fig 2 was carried out using publicly available datasets. TCGA data was obtained from https://xenabrowser.net/datapages/. Different EMT scoring metrics were calculated as described previously [44].

The set of differential equations used to simulate the dynamics of the EMT / MET regulatory circuit are enumerated in the SI. The SI also includes all the model parameters and a description of how epigenetic feedback was incorporated into the mathematical model of EMT / MET regulation. Finally, the analysis shown in Fig 7 was carried out using the population-level model described previously (3). The computer code used to generate the data shown in Fig 7 is available online (https://github.com/st35/cancer-EMT-heterogeneity-noise/tree/master/ExternalIntervention; see files DailyIntervention_MET.cpp and DailyIntervention_GRHL2_MET.cpp).

## Author contributions

MKJ designed research; MKJ and HL supervised research; WJ, ST, PC, AC conducted research; AR, MKJ, HL analyzed data; all authors participated in writing and revision of the manuscript.

## Notes

### Competing Interest Statement

The authors have declared no competing interest.

## References

1. Nieto MA, Huang RY, Jackson RA, Thiery JP. Emt. Cell. 2016; 166: 21–45.

2. Jolly MK, Mani SA, Levine H. Hybrid epithelial/mesenchymal phenotype(s): The ‘fittest’ for metastasis? Biochim Biophys Acta - Rev Cancer. 2018; 1870: 151–7. doi: 10.1016/j.bbcan.2018.07.001.

3. Tripathi S, Chakraborty P, Levine H, Jolly MK. A mechanism for epithelial-mesenchymal heterogeneity in a population of cancer cells. PLoS Comput Biol. 2020; 16: e1007619. doi: 10.1101/592691.

4. Pastushenko I, Brisebarre A, Sifrim A, Fioramonti M, Revenco T, Boumahdi S, Van Keymeulen A, Brown D, Moers V, Lemaire S, De Clercq S, Minguijón E, Balsat C, et al. Identification of the tumour transition states occurring during EMT. Nature. 2018; 556: 463–8. doi: 10.1038/s41586-018-0040-3.

5. Goetz H, Melendez-Alvarez JR, Chen L, Tian X-J. A plausible accelerating function of intermediate states in cancer metastasis. PLOS Comput Biol [Internet]. 2020; 16: e1007682. doi: 10.1371/journal.pcbi.1007682.

6. Huang RY-J, Wong MK, Tan TZ, Kuay KT, Ng a HC, Chung VY, Chu Y-S, Matsumura N, Lai H-C, Lee YF, Sim W-J, Chai C, Pietschmann E, et al. An EMT spectrum defines an anoikis-resistant and spheroidogenic intermediate mesenchymal state that is sensitive to e-cadherin restoration by a src-kinase inhibitor, saracatinib (AZD0530). Cell Death Dis. 2013; 4: e915. doi: 10.1038/cddis.2013.442.

7. Tripathi SC, Peters HL, Taguchi A, Katayama H, Wang H, Momin A, Jolly MK, Celiktas M, Rodriguez-Canales J, Liu H, Behrens C, Wistuba II, Ben-Jacob E, et al. Immunoproteasome deficiency is a feature of non-small cell lung cancer with a mesenchymal phenotype and is associated with a poor outcome. Proc Natl Acad Sci U S A. 2016; 113. doi: 10.1073/pnas.1521812113.

8. Ocaña OH, Córcoles R, Fabra A, Moreno-Bueno G, Acloque H, Vega S, Barrallo-Gimeno A, Cano A, Nieto MA. Metastatic colonization requires the repression of the epithelial-mesenchymal transition inducer Prrx1. Cancer Cell. 2012; 22: 709–24. doi: 10.1016/j.ccr.2012.10.012.

9. Tsai JH, Donaher JL, Murphy DA, Chau S, Yang J. Spatiotemporal regulation of epithelial-mesenchymal transition is essential for squamous cell carcinoma metastasis. Cancer Cell. 2012; 22: 725–36. doi: 10.1016/j.ccr.2012.09.022.

10. Hari K, Sabuwala B, Subramani BV, Porta C La, Zapperi S, Font-Clos F, Jolly MK. Identifying inhibitors of epithelial-mesenchymal plasticity using a network topology based approach. bioRxiv. Cold Spring Harbor Laboratory; 2019;: 854307. doi: 10.1101/854307.

11. Katsuno Y, Meyer DS, Zhang Z, Shokat KM, Akhurst RJ, Miyazono K, Dernyck R. Chronic TGF-β exposure drives stabilized EMT, tumor stemness, and cancer drug resistance with vulnerability to bitopic mTOR inhibition. Sci Signal. 2019; 12: eaau8544. doi: 10.1126/scisignal.aau8544.

12. Gregory PA, Bracken CP, Smith E, Bert AG, Wright J a, Roslan S, Morris M, Wyatt L, Farshid G, Lim Y-Y, Lindeman GJ, Shannon MF, Drew P a, et al. An autocrine TGF-beta/ZEB/miR-200 signaling network regulates establishment and maintenance of epithelial-mesenchymal transition. Mol Biol Cell. 2011; 22: 1686–98. doi: 10.1091/mbc.E11-02-0103.

13. Jia W, Deshmukh A, Mani SA, Jolly MK, Levine H. A possible role for epigenetic feedback regulation in the dynamics of the Epithelial-Mesenchymal Transition (EMT). Phys Biol. 2019; 16: 066004. doi: 10.1101/651620.

14. Eichelberger L, Saini M, Moreno HD, Klein C, Bartsch JM, Falcone M, Reitberger M, Espinet E, Vogel V, Graf E, Schwarzmayr T, Strom T-M, Lehmann M, et al. Maintenance of epithelial traits and resistance to mesenchymal reprogramming promote proliferation in metastatic breast cancer. bioRxiv. 2020;: 998823. doi: 10.1101/2020.03.19.998823.

15. Richardson LS, Taylor RN, Menon R. Reversible EMT and MET mediate amnion remodeling during pregnancy and labor. Sci Signal. 2020; 13: e11y1486. doi: 10.1126/scisignal.aay1486.

16. Shinde A, Hardy SD, Kim D, Akhand SS, Jolly MK, Wang WH, Anderson JC, Khodadadi RB, Brown WS, George JT, Liu S, Wan J, Levine H, et al. Spleen tyrosine kinase– mediated autophagy is required for epithelial–mesenchymal plasticity and metastasis in breast cancer. Cancer Res. 2019; 79: 1831–43. doi: 10.1158/0008-5472.CAN-18-2636.

17. Zhou X, Wang J, Chen J, Qi Y, Di Nan, Jin L, Qian X, Wang X, Chen Q, Liu X, Xu Y. Optogenetic control of epithelial-mesenchymal transition in cancer cells. Sci Rep. 2018; 8: 14098. doi: 10.1038/s41598-018-32539-3.

18. Tian X-J, Zhang H, Xing J. Coupled Reversible and Irreversible Bistable Switches Underlying TGFβ-induced Epithelial to Mesenchymal Transition. Biophys J. 2013; 105: 1079–89. doi: 10.1016/j.bpj.2013.07.011.

19. Zhang J, Tian X-J, Zhang H, Teng Y, Li R, Bai F, Elankumaran S, Xing J. TGF-β–induced epithelial-to-mesenchymal transition proceeds through stepwise activation of multiple feedback loops. Sci Signal. 2014; 7: ra91. doi: 10.1126/scisignal.2005304.

20. Watanabe K, Panchy N, Noguchi S, Suzuki H, Hong T. Combinatorial perturbation analysis reveals divergent regulations of mesenchymal genes during epithelial-to-mesenchymal transition. npj Syst Biol Appl. 2019; 5: 21. doi: 10.1038/s41540-019-0097-0.

21. Steinway SN, Zañudo JGT, Michel PJ, Feith DJ, Loughran TP, Albert R. Combinatorial interventions inhibit TGFβ-driven epithelial-to-mesenchymal transition and support hybrid cellular phenotypes. npj Syst Biol Appl. Nature Publishing Group; 2015; 1: 15014. doi: 10.1038/npjsba.2015.14.

22. Somarelli JA, Shelter S, Jolly MK, Wang X, Bartholf Dewitt S, Hish AJ, Gilja S, Eward WC, Ware KE, Levine H, Armstrong AJ, Garcia-Blanco MA. Mesenchymal-epithelial transition in sarcomas is controlled by the combinatorial expression of miR-200s and GRHL2. Mol Cell Biol. 2016; 36: 2503–13. doi: 10.1128/MCB.00373-16.

23. Nihan Kilinc A, Sugiyama N, Reddy Kalathur RK, Antoniadis H, Birogul H, Ishay-Ronen D, George JT, Levine H, Kumar Jolly M, Christofori G. Histone deacetylases, Mbd3/NuRD, and Tet2 hydroxylase are crucial regulators of epithelial–mesenchymal plasticity and tumor metastasis. Oncogene. Nature Publishing Group; 2019; 39: 1498–513. doi: 10.1038/s41388-019-1081-2.

24. Brown KA, Aakre ME, Gorska AE, Price JO, Eltom SE, Pietenpol JA, Moses HL. Induction by transforming growth factor-β1 of epithelial to mesenchymal transition is a rare event in vitro. Breast Cancer Res. 2004; 6: R215–31. doi: 10.1186/bcr778.

25. Cieply B, Riley IV P, Pifer PM, Widmeyer J, Addison JB, Ivanov A V., Denvir J, Frisch SM. Suppression of the epithelial-mesenchymal transition by grainyhead-like-2. Cancer Res. 2012; 72: 2440–53. doi: 10.1158/0008-5472.CAN-11-4038.

26. Cieply B, Farris J, Denvir J, Ford HL, Frisch SM. Epithelial-Mesenchymal Transition and Tumor Suppression Are Controlled by a Reciprocal Feedback Loop between ZEB1 and Grainyhead-like-2. Cancer Res. 2013; 73: 6299–309. doi: 10.1158/0008-5472.CAN-12-4082.

27. Xiang X, Deng Z, Zhuang X, Ju S, Mu J, Jiang H, Zhang L, Yan J, Miller D, Zhang HG. Grhl2 Determines the Epithelial Phenotype of Breast Cancers and Promotes Tumor Progression. PLoS One. 2012; 7: e50781. doi: 10.1371/journal.pone.0050781.

28. Huh D, Paulsson J. Non-genetic heterogeneity from stochastic partitioning at cell division. Nat Genet. 2011; 43: 95–100. doi: 10.1038/ng.729.

29. Huh D, Paulsson J. Random partitioning of molecules at cell division. Proc Natl Acad Sci U S A. 2011; 108: 15004–9. doi: 10.1073/pnas.1013171108.

30. Soltani M, Vargas-Garcia CA, Antunes D, Singh A. Intercellular Variability in Protein Levels from Stochastic Expression and Noisy Cell Cycle Processes. PLoS Comput Biol. 2016; 12: e1004972. doi: 10.1371/journal.pcbi.1004972.

31. Drápela S, Bouchal J, Jolly MK, Culig Z. ZEB1: A Critical Regulator of Cell Plasticity, DNA Damage Response, and Therapy Resistance. Front Mol Biosci. 2020; 7: 36. doi: 10.3389/fmolb.2020.00036.

32. Miyamoto T, Furusawa C, Kaneko K. Pluripotency, Differentiation, and Reprogramming: A Gene Expression Dynamics Model with Epigenetic Feedback Regulation. PLoS Comput Biol. 2015; 11: e1004476. doi: 10.1371/journal.pcbi.1004476.

33. Matshushita Y, Kaneko K. Homeorhesis in Waddington’s landscape by epigenetic feedback regulation. Phys Rev Res. 2020; 2: 023083. doi: 10.1103/PhysRevResearch.2.023083.

34. Jolly MK, Huang B, Lu M, Mani SA, Levine H, Ben-Jacob E. Towards elucidating the connection between epithelial − mesenchymal transitions and stemness. J R Soc Interface. 2014; 11: 20140962. doi: 10.1098/rsif.2014.0962.

35. Grosse-Wilde A, Fouquier d’ Herouei A, McIntosh E, Ertaylan G, Skupin A, Kuestner RE, del Sol A, Walters K-A, Huang S. Stemness of the hybrid epithelial/mesenchymal state in breast cancer and its association with poor survival. PLoS One. 2015; 10: e0126522. doi: 10.1371/journal.pone.0126522.

36. Jolly MK, Tripathi SC, Jia D, Mooney SM, Celiktas M, Hanash SM, Mani SA, Pienta KJ, Ben-Jacob E, Levine H. Stability of the hybrid epithelial/mesenchymal phenotype. Oncotarget. 2016; 7: 27067–84. doi: 10.18632/oncotarget.8166.

37. Quan Y, Jin R, Huang A, Zhao H, Feng B, Zang L, Zheng M. Downregulation of GRHL2 inhibits the proliferation of colorectal cancer cells by targeting ZEB1. Cancer Biol Ther. 2014; 15: 878–87. doi: 10.4161/cbt.28877.

38. Chung VY, Tan TZ, Tan M, Wong MK, Kuay KT, Yang Z, Ye J, Muller J, Koh CM, Guccione E, Thiery JP, Huang RY-J. GRHL2-miR-200-ZEB1 maintains the epithelial status of ovarian cancer through transcriptional regulation and histone modification. Sci Rep. Nature Publishing Group; 2016; 6: 19943. doi: 10.1038/srep19943.

39. Chung VY, Tan TZ, Ye J, Huang R-L, Lai H-C, Kappei D, Wollmann H, Guccione E, Huang RY-J. The role of GRHL2 and epigenetic remodeling in epithelial–mesenchymal plasticity in ovarian cancer cells. Commun Biol [Internet]. Springer US; 2019; 2: 272. doi: 10.1038/s42003-019-0506-3.

40. Xiang J, Fu X, Ran W, Wang Z. Grhl2 reduces invasion and migration through inhibition of TGFβ-induced EMT in gastric cancer. Oncogenesis. 2017; 6: e284. doi: 10.1038/oncsis.2016.83.

41. Chen W, Yi JK, Shimane T, Mehrazarin S, Lin YL, Shin KH, Kim RH, Park NH, Kang MK. Grainyhead-like 2 regulates epithelial plasticity and stemness in oral cancer cells. Carcinogenesis. 2016; 37: 500–10. doi: 10.1093/carcin/bgw027.

42. Nishino H, Takano S, Yoshitomi H, Suzuki K, Kagawa S, Shimazaki R, Shimizu H, Furukawa K, Miyazaki M, Ohtsuka M. Grainyhead-like 2 (GRHL2) regulates epithelial plasticity in pancreatic cancer progression. Cancer Med. 2017; 6: 2686–96. doi: 10.1002/cam4.1212.

43. Barretina J, Caponigro G, Stransky N, Venkatesan K, Margolin A a., Kim S, Wilson CJ, Lehár J, Kryukov G V., Sonkin D, Reddy A, Liu M, Murray L, et al. The Cancer Cell Line Encyclopedia enables predictive modelling of anticancer drug sensitivity. Nature. 2012; 483: 603–7. doi: 10.1038/nature11003.

44. Chakraborty P, George JT, Tripathi S, Levine H, Jolly MK. Comparative study of transcriptomics-based scoring metrics for the epithelial-hybrid-mesenchymal spectrum. Front Bioeng Biotechnol. 2020; 8: 220. doi: 10.3389/fbioe.2020.00220.

45. Wang Y, Shi J, Chai K, Ying X, Zhou B. The Role of Snail in EMT and Tumorigenesis. Curr Cancer Drug Targets. 2013; 13: 963–72. doi: 10.2174/15680096113136660102.

46. Frisch SM, Farris JC, Pifer PM. Roles of Grainyhead-like transcription factors in cancer. Oncogene [Internet]. Nature Publishing Group; 2017; 36: 6067–73. doi: 10.1038/onc.2017.178.

47. Panchy N, Azeredo-Tseng C, Luo M, Randall N, Hong T. Integrative Transcriptomic Analysis Reveals a Multiphasic Epithelial–Mesenchymal Spectrum in Cancer and Non-tumorigenic Cells. Front Oncol. Frontiers Media S.A.; 2020; 9: 1479. doi: 10.3389/fonc.2019.01479.

48. Varma S, Cao Y, Tagne JB, Lakshminarayanan M, Li J, Friedman TB, Morell RJ, Warburton D, Kotton DN, Ramirez MI. The transcription factors grainyhead-like 2 and NK2-homeobox 1 form a regulatory loop that coordinates lung epithelial cell morphogenesis and differentiation. J Biol Chem. 2012; 287: 37282–95. doi: 10.1074/jbc.M112.408401.

49. Tripathi S, Levine H, Jolly MK. The Physics of Cellular Decision-Making during Epithelial-Mesenchymal Transition. Annu Rev Biophys. 2020; 49: 1–18. doi: 10.1146/annurev-biophys-121219-081557.

50. Milo R, Jorgensen P, Moran U, Weber G, Springer M. BioNumbers--the database of key numbers in molecular and cell biology. Nucleic Acids Res. 2010; 38: D750–3. doi: 10.1093/nar/gkp889.

51. Karacosta LG, Anchang B, Ignatiadis N, Kimmey SC, Benson JA, Shrager JB, Tibshirani R, Bendall SC, Plevritis SK. Mapping Lung Cancer Epithelial-Mesenchymal Transition States and Trajectories with Single-Cell Resolution. Nat Commun. 2019; 10: 5587. doi: 10.1101/570341.

52. Celià-Terrassa T, Bastian C, Liu DD, Ell B, Aiello NM, Wei Y, Zamalloa J, Blanco AM, Hang X, Kunisky D, Li W, Williams ED, Rabitz H, et al. Hysteresis control of epithelial-mesenchymal transition dynamics conveys a distinct program with enhanced metastatic ability. Nat Commun. Springer US; 2018; 9: 5005. doi: 10.1038/s41467-018-07538-7.

53. Devaraj V, Bose B. Morphological State Transition Dynamics in EGF-Induced Epithelial to Mesenchymal Transition. J Clin Med. 2019; 8: 911. doi: 10.3390/jcm8070911.

54. Stylianou N, Lehman ML, Wang C, Fard AT, Rockstroh A, Fazli L, Jovanovic L, Ward M, Sadowski MC, Kashyap AS, Buttyan R, Gleave ME, Westbrook TF, et al. A molecular portrait of epithelial–mesenchymal plasticity in prostate cancer associated with clinical outcome. Oncogene. 2018;: Epub ahead of print. doi: 10.1038/s41388-018-0488-5.

55. Xu S, Ware KE, Ding Y, Kim S-Y, Sheth M, Rao S, Chan W, Armstrong AJ, Eward WC, Jolly M, Somarelli JA. An integrative systems biology and experimental approach identifies convergence of epithelial plasticity, metabolism, and autophagy to promote chemoresistance. bioRxiv. 2018;. doi: 10.1101/365833.

56. Mandal M, Ghosh B, Anura A, Mitra P, Pathak T, Chatterjee J. Modeling continuum of epithelial mesenchymal transition plasticity. Integr Biol [Internet]. Royal Society of Chemistry; 2016; 8: 167–76. doi: 10.1039/C5IB00219B.

57. Mehrazarin S, Chen W, Oh J-E, Liu ZX, Kang KL, Yi JK, Kim RH, Shin K-H, Park N-H, Kang MK. p63 Gene is Regulated by Grainyhead-Like 2 (GRHL2) Through Reciprocal Feedback and Determines Epithelial Phenotype in Human Keratinocytes. J Biol Chem. 2015; 290: 19999–20008. doi: 10.1074/jbc.M115.659144.

58. Jolly MK, Boareto M, Debeb BG, Aceto N, Farach-Carson MC, Woodward WA, Levine H. Inflammatory Breast Cancer: a model for investigating cluster-based dissemination. NPJ Breast Cancer. 2017; 3: 21. doi: https://doi.org/10.1101/119479.

59. Aue A, Hinze C, Walentin K, Ruffert J, Yurtdas Y, Werth M, Chen W, Rabien A, Kilic E, Schulzke J-D, Schumann M, Schmidt-Ott KM. A Grainyhead-Like 2/Ovo-Like 2 Pathway Regulates Renal Epithelial Barrier Function and Lumen Expansion. J Am Soc Nephrol. 2015; 26: 2704–15. doi: 10.1681/ASN.2014080759.

60. Roca H, Hernandez J, Weidner S, McEachin RC, Fuller D, Sud S, Schumann T, Wilkinson JE, Zaslavsky A, Li H, Maher CA, Daignault-Newton S, Healy PN, et al. Transcription Factors OVOL1 and OVOL2 Induce the Mesenchymal to Epithelial Transition in Human Cancer. PLoS One. 2013; 8: e76773. doi: 10.1371/journal.pone.0076773.

61. Somarelli JA, Shetler S, Jolly MK, Wang X, Dewitt SB, Hish AJ, Gilja S, Eward WC, Ware KE, Levine H, others. Mesenchymal-Epithelial Transition in Sarcomas Is Controlled by the Combinatorial Expression of MicroRNA 200s and GRHL2. Mol Cell Biol. Am Soc Microbiol; 2016; 36: 2503–13.

62. Tam WL, Weinberg RA. The epigenetics of epithelial-mesenchymal plasticity in cancer. Nat Med. 2013; 19: 1438–49. doi: 10.1038/nm.3336.

63. Jacobs J, Atkins M, Davie K, Imrichova H, Romanelli L, Christiaens V, Hulselmans G, Potier D, Wouters J, Taskiran II, Paciello G, González-Blas CB, Koldere D, et al. The transcription factor Grainy head primes epithelial enhancers for spatiotemporal activation by displacing nucleosomes. Nat Genet. 2018; 50: 1011–20. doi: 10.1038/s41588-018-0140-x.

64. Chen AF, Liu AJ, Krishnakumar R, Freimer JW, DeVeale B, Blelloch R. GRHL2-Dependent Enhancer Switching Maintains a Pluripotent Stem Cell Transcriptional Subnetwork after Exit from Naive Pluripotency. Cell Stem Cell. 2018; 23: 226–38. doi: 10.1016/j.stem.2018.06.005.

65. Jolly MK, Kulkarni P, Weninger K, Orban J, Levine H. Phenotypic Plasticity, Bet-Hedging, and Androgen Independence in Prostate Cancer: Role of Non-Genetic Heterogeneity. Front Oncol. 2018; 8: 1–12. doi: 10.3389/fonc.2018.00050.

66. Ozbudak EM, Thattai M, Lim HN, Shraiman BI, van Oudenaarden A. Multistability in the lactose utilization network of Escherichia coli. Nature. 2004; 427: 737–40. doi: 10.1038/nature02298.

67. Jolly MK, Tripathi SC, Somarelli JA, Hanash SM, Levine H. Epithelial/mesenchymal plasticity: how have quantitative mathematical models helped improve our understanding? Mol Oncol. 2017; 11: 739–54. doi: 10.1002/1878-0261.12084.

68. Jolly MK, Jia D, Boareto M, Mani SA, Pienta KJ, Ben-Jacob E, Levine H. Coupling the modules of EMT and stemness: A tunable “stemness window” model. Oncotarget. 2015; 6: 25161–74. doi: 10.18632/oncotarget.4629.

69. Youssef G, Gammon L, Ambler L, Wicker B, Patel S, Cottom H, Piper K, Mackenzie IC, Philpott MP, Biddle A. Disseminating cells in human tumous acquire an EMT stem cell state that is predictive of metastasis. bioRxiv. 2020;: 029009. doi: 10.1101/2020.04.07.029009.

70. Kröger C, Afeyan A, Mraz J, Eaton EN, Reinhardt F, Khodor YL, Thiru P, Bierie B, Ye X, Burge CB, Weinberg RA. Acquisition of a hybrid E/M state is essential for tumorigenicity of basal breast cancer cells. Proc Natl Acad Sci. 2019; 116: 7353–62. doi: 10.1073/pnas.1812876116.

71. Bierie B, Pierce SE, Kroeger C, Stover DG, Pattabiraman DR, Thiru P, Liu Donaher J, Reinhardt F, Chaffer CL, Keckesova Z, Weinberg RA. Integrin-β4 identifies cancer stem cell-enriched populations of partially mesenchymal carcinoma cells. Proc Natl Acad Sci [Internet]. 2017; 114: E2337–2346. doi: 10.1073/pnas.1618298114.

72. Celià-Terrassa T, Meca-Cortés Ó, Mateo F, De Paz AM, Rubio N, Arnal-Estapé A, Ell BJ, Bermudo R, Díaz A, Guerra-Rebollo M, Lozano JJ, Estarás C, Ulloa C, et al. Epithelial-mesenchymal transition can suppress major attributes of human epithelial tumor-initiating cells. J Clin Invest. 2012; 122: 1849–68. doi: 10.1172/JCI59218.

73. Liu S, Cong Y, Wang D, Sun Y, Deng L, Liu Y, Martin-Trevino R, Shang L, McDermott SP, Landis MD, Hong S, Adams A, D’Angelo R, et al. Breast cancer stem cells transition between epithelial and mesenchymal states reflective of their normal counterparts. Stem Cell Reports. 2014; 2: 78–91. doi: 10.1016/j.stemcr.2013.11.009.

74. Bocci F, Gearhart-Serna L, Boareto M, Ribeiro M, Ben-Jacob E, Devi GR, Levine H, Onuchic JN, Jolly MK. Toward understanding cancer stem cell heterogeneity in the tumor microenvironment. Proc Natl Acad Sci U S A. 2019; 116: 148–57. doi: 10.1073/pnas.1815345116.

75. Beerling E, Seinstra D, de Wit E, Kester L, van der Velden D, Maynard C, Schäfer R, van Diest P, Voest E, van Oudenaarden A, Vrisekoop N, van Rheenen J. Plasticity between Epithelial and Mesenchymal States Unlinks EMT from Metastasis-Enhancing Stem Cell Capacity. Cell Rep. 2016; 14: 2281–8. doi: 10.1016/j.celrp.201602.034.

76. Lovisa S, LeBleu VS, Tampe B, Sugimoto H, Vadnagara K, Carstens JL, Wu C-C, Hagos Y, Burckhardt BC, Pentcheva-Hoang T, Nischal H, Allison JP, Zeisberg M, et al. Epithelial-to-mesenchymal transition induces cell cycle arrest and parenchymal damage in renal fibrosis. Nat Med. 2015; 21: 998–1009. doi: 10.1038/nm.3902.

77. Vega S, Morales A V., Ocaña OH, Valdés F, Fabregat I, Nieto MA. Snail blocks the cell cycle and confers resistance to cell death. Genes Dev. 2004; 18: 1131–43. doi: 10.1101/gad.294104.

78. Vittadello ST, McCue SW, Gunasingh G, Haass NK, Simpson MJ. Examining go-or-grow using flourescent cell-cycle indicators and cell-cycle-inhibiting drugs. Biophys J. 2020; 118: 1243–7. doi: 10.1016/j.bpj.2020.01.036.

