## Supplementary Information for "Epigenetic feedback and stochastic partitioning during cell division can drive resistance to EMT"

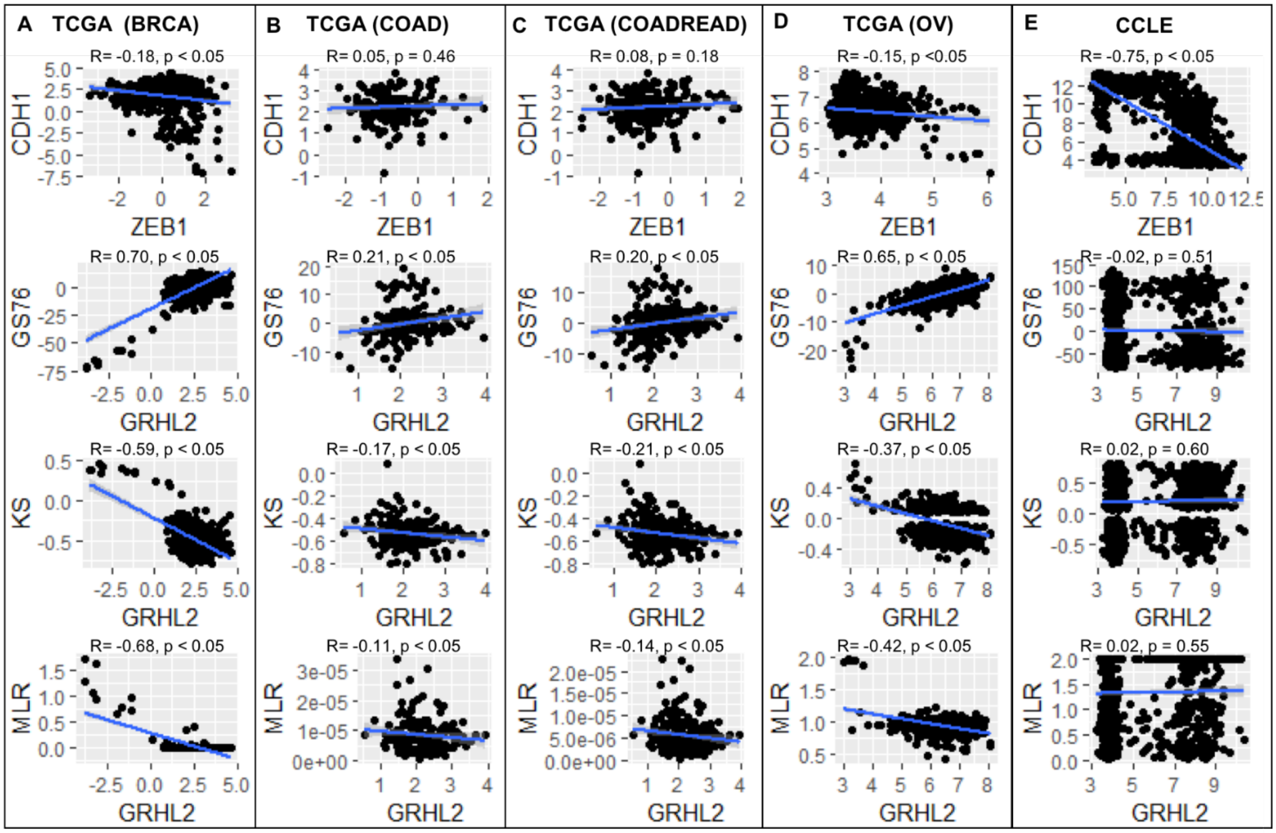


***Figure S1: GRHL2 correlates with an epithelial phenotype.*** *Scatter plots showing correlation of GRHL2 with EMT scores calculated via three metrics (GS76, KS, MLR), and that of ZEB1 vs. CDH1 in TCGA datasets and CCLE – A) breast cancer, B) colon adenocarcinoma, C) colorectal adenocarcinoma, D) ovarian carcinoma, E) CCLE. R, p denote Spearman's correlation coefficient and corresponding p-value for corresponding plot.*

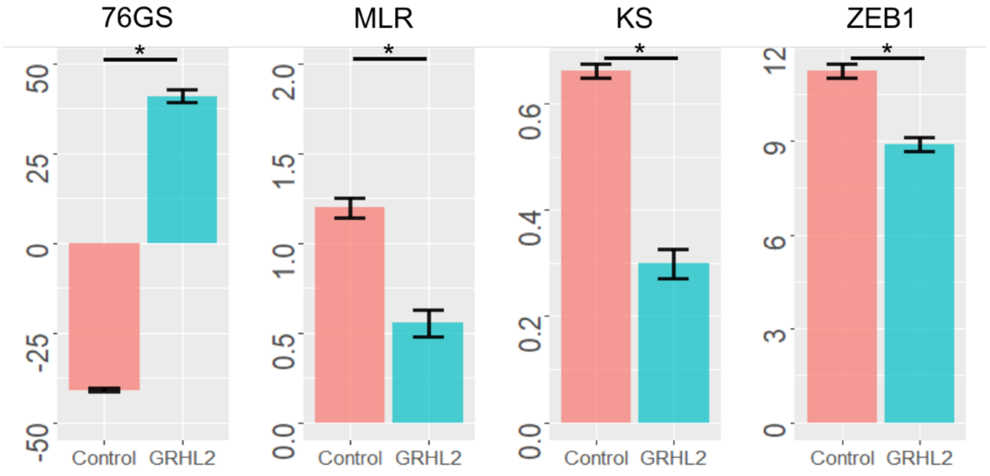

***Figure S2: GRHL2 correlates with EMT scores.*** *Bar plots showing the mean EMT scores calculated via 76GS, KS and MLR scoring metric for HMLE+Twist-ER cells expressing GRHL2/pMIG ('GRHL2') vs. HMLE+Twist-ER cells expressing empty pMIG ('control'). The last panel shows corresponding ZEB1 levels (GSE36081). Error bars show standard deviation; n=3.*

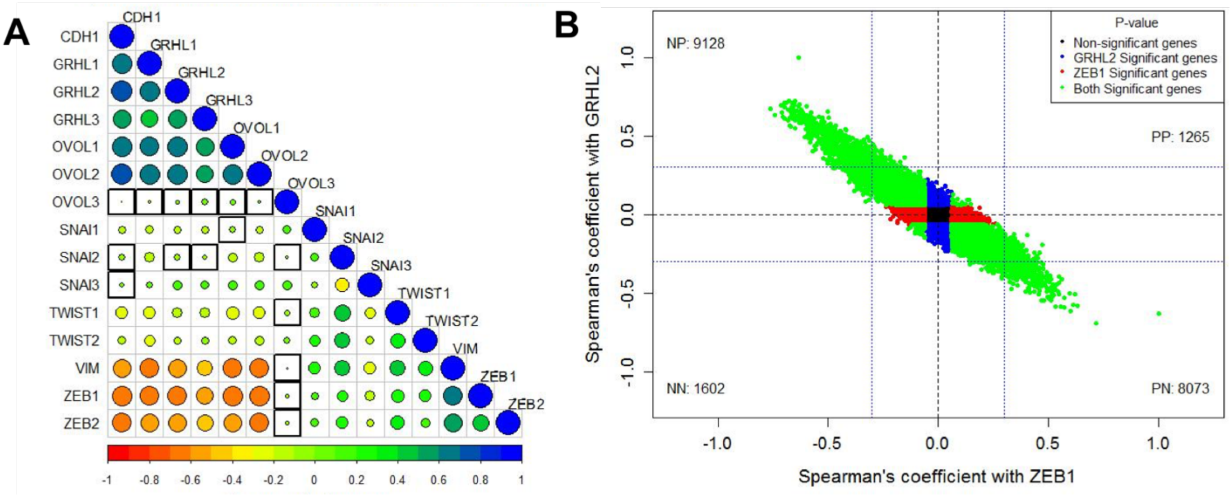


***Figure S3: GRHL2/ZEB1 axis correlates with EMT/MET across cancer types***. *A) Pairwise Spearman's correlation between different EMT and MET regulatory genes in the CCLE dataset. Spearman's correlation value (cor) of each gene pair is represented as the size of the circle and filled with corresponding color from the color palette represented below the ranging from -1(red) to +1(blue). Boxes highlighted by the black squares represent insignificant (p>0.01) correlation. B) Scatter plot of genes correlated using Pearson correlation method with GRHL2 and ZEB1 in CCLE dataset. Each dot represents one gene and coordinates are Spearson's cor values with ZEB1 and GRHL2. Color of the dots is based on the p-value obtained from correlation test, Blue dots for genes having p<0.05 with GRHL2 and p>0.05 with ZEB1, Red dots for genes having p<0.05 with ZEB1 and p>0.05 with GRHL2, Green dots for genes having p<0.05 with GRHL2 and ZEB1 and Black dots for genes having p>0.05 05 with GRHL2 and ZEB1. Numbers in each quadrant represent the number of genes in that quadrant.*

### 1. Theoretical model for EMT

The core EMT network consists of two mutually inhibiting loop: miR-200/ZEB and miR-34/SNAIL. Here, GRHL2 forms another mutually inhibiting loop with ZEB (1, 2). Deterministic equations for miR-200/ZEB circuit with the external signal as SNAIL (3,4) are given below:

$$\dot{\mu}_{200} = g_{\mu_{200}} H^s(Z, \lambda_{z,\mu_{200}}) H^s(S, \lambda_{S,\mu_{200}}) - m_z Y_\mu(\mu_{200}) - k_\mu \mu_{200}$$

$$\dot{m}_z = g_{m_z} H^s(Z, \lambda_{z,m_z}) H^s(S, \lambda_{s,m_z}) H^s(G, \lambda_{G,m_z}) - m_z Y_m(\mu_{200}) - k_{m_z} m_z$$

$$\dot{Z} = g_z m_Z L(\mu_{200}) - k_z Z$$

and those for miR-34/SNAIL circuit with I as an external signal are:

$$\dot{\mu}_{34} = g_{\mu_{34}} H^s(S, \lambda_{s,\mu_{34}}) H^s(Z, \lambda_{z,\mu_{34}}) - m_s Y_\mu(\mu_{34}) - k_{\mu_{34}} \mu_{34}$$

$$\dot{m}_s = g_{m_s} H^s(S, \lambda_{s,m_s}) H^s(I, \lambda_{I,m_s}) - m_s Y_m(\mu_{34}) - k_{m_s} m_s$$

$$\dot{S} = g_s m_s L(\mu_{34}) - k_s S$$

and for the GRHL2/ZEB loop:

$$\dot{m}_G = g_{m_G} H^s(Z, \lambda_{Z,m_G}) H^s(G, \lambda_{G,m_G}) - k_{m_G} m_G$$

$$\dot{G} = g_G m_G - k_G G$$

where $g$ is the innate synthesis rate for corresponding microRNA/mRNA/protein, $k$ is the corresponding innate degradation rate. Here $H^S$ represents the shifted Hill function which is defined as:

$$H^s(B) = \frac{1 + \lambda \left(\frac{B}{B_0}\right)^{n_B}}{1 + \left(\frac{B}{B_0}\right)^{n_B}}$$

where $\lambda$ is the fold change regulated by protein B. $\lambda$>1 for activation and $\lambda$<1 for inhibition.

The external signal I that we use here can be written as the stochastic differential equation:
$\dot{I} = \beta(I_0 - I) + \eta(t)$
where $\eta(t)$ satisfies the condition that $< \eta(t), n(t') \geq \Gamma\delta(t - t')$. Here $I_0$ is set at 50 K molecules, $\beta$ as 0.04 hour$^{-1}$, and $\Gamma$ as 1000 (K molecules/hour)$^2$.
The initial value of I is fixed to lie at the middle of the tristable region {E, E/M, M}.

**2.Theoretical model for microRNA/TF (transcription factor) circuits**

In microRNA-based chimeric (MBC) circuits, microRNA(miR) molecules bind to the 3' UTR of the corresponding mRNA of the target protein in order to form a new miR-mRNA complex. In this way, miRs can inhibit the translation of mRNA and/or active mRNA degradation, and also can be degraded or recycled themselves (5). Based on the different miR-mRNA complexes by binding/unbinding chemical reactions, a new computational model has been established (6).

Assume an mRNA has $n$ miR binding sites, so there are $n + 1$ possible configurations of mRNA. If $i$ is used to represent the number of miRs binding with one mRNA, the value of $i$ can be from 0 to $n$. The binding process can be considered independent among different binding sites because a miR is 22 nt long and it recognizes mRNA by a seed sequence only of 7-8 nt. Assuming that

the binding/unbinding rate of miR and mRNA is much faster than the molecule production/degradation rate, at equilibrium, the concentration of mRNA $[m_i]$ where $i$ miR molecules bind to i binding sites of mRNA satisfies $r_{\mu+}\mu[m_i] = r_{\mu-}[m_{i+1}]$.Here $r_{\mu+}$ is the binding rate, $r_{\mu-}$ is the unbinding rate, and $\mu$ is the miR concentration. If we set $\mu_0 = r_{\mu-}/r_{\mu+}$, then we get $[m_i] = \left(\frac{\mu}{\mu_0}\right)^i [m_0]$. The total mRNA concentration is $m = \sum_{i=0}^{n} C_n^i [m_i]$. The second equation can be rewritten as $[m_i] = mM_n^i(\mu)$, while $M_n^i(\mu) = \frac{\left(\frac{\mu}{\mu_0}\right)^i}{\left(1+\frac{\mu}{\mu_0}\right)^n}$. Finally, we can write the total translation rate $\sum_{i=0}^{n} l_i C_n^i [m_i] = m\sum_{i=0}^{n} l_i C_n^i M_n^i(\mu) = mL(\mu)$ , the total mRNA active degradation rate $\sum_{i=0}^{n} \gamma_{mi} C_n^i [m_i] = m\sum_{i=0}^{n} \gamma_{mi} C_n^i M_n^i(\mu) = mY_m(\mu)$ and the total miR active degradation rate $\sum_{i=0}^{n} \gamma_{\mu i} C_n^i [m_i] = m\sum_{i=0}^{n} \gamma_{\mu i} C_n^i M_n^i(\mu) = mY_\mu(\mu)$.
Here, $l_i$ is the individual translation rate, $\gamma_{mi}$ is the individual active degradation rate of mRNA and $\gamma_{\mu i}$ is the individual active degradation rate of miR. This model can capture different mechanisms by choosing different parameters.

#### 3. Epigenetic feedback regulation term

In the EMT model, we tested epigenetic feedback through two different pathways. The dynamic equation of epigenetic feedback on GRHL2's self-activation is:

$$\dot{G}_{m_G}^0 = \frac{G_{m_G}^0(0) - G_{m_G}^0 - \alpha G}{\zeta}$$

Similarly, an epigenetic feedback on ZEB's inhibition from GRHL2 is modeled via:

$$G_{m_Z}^0 = \frac{G_{m_Z}^0(0) - G_{m_Z}^0 - \alpha G}{\zeta}$$

where $\zeta$ is a timescale factor and chosen to be 100 (hours). $\alpha$ represents the strength of epigenetic feedback. Larger $\alpha$ corresponds to stronger epigenetic feedback. $\alpha$ has an upper bound (usually between 0.01-0.25) because of the restriction that the numbers of all molecules must be positive. For GRHL2's self-activation, high level of GRHL2 can activate the expression of GRHL2 itself. Meanwhile, for the inhibition of ZEB by GRHL2, high levels of GRHL2 can suppress the production of ZEB.

#### 4.Parameters for the EMT model

**Table SI 1. List of parameters used in shifted Hill functions**

| Description | Fold change | Value | # of binding sites | Value | Threshold | Value (K molecules) |
|---|---|---|---|---|---|---|
| **Inhibition on miR-200 by ZEB** | $\lambda_{Z,\mu_{200}}$ | 0.1 | $n_{Z,\mu_{200}}$ | 3 | $Z_{\mu_{200}}^0$ | 220 |
| **Inhibition on miR-200 by SNAIL** | $\lambda_{S,\mu_{200}}$ | 0.1 | $n_{S,\mu_{200}}$ | 2 | $S_{\mu_{200}}^0$ | 180 |

| | | | | | | |
|---|---|---|---|---|---|---|
| **Self-activation of ZEB** | $\lambda_{Z,m_z}$ | 7.5 | $n_{Z,m_z}$ | 2 | $Z^0_{m_z}$ | 25 |
| **Activation on ZEB by SNAIL** | $\lambda_{S,m_z}$ | 10.0 | $n_{S,m_z}$ | 2 | $S^0_{m_z}$ | 180 |
| **Inhibition on ZEB by GRHL2** | $\lambda_{G,m_z}$ | 0.65 | $n_{G,m_z}$ | 1 | $G^0_{m_z}$ | 25 |
| **Inhibition on miR-34 by SNAIL** | $\lambda_{S,\mu_{34}}$ | 0.1 | $n_{S,\mu_{34}}$ | 1 | $S^0_{\mu_{34}}$ | 300 |
| **Inhibition on miR-34 by ZEB** | $\lambda_{Z,\mu_{34}}$ | 0.2 | $n_{Z,\mu_{34}}$ | 2 | $Z^0_{\mu_{34}}$ | 600 |
| **Self-inhibition of SNAIL** | $\lambda_{S,m_s}$ | 0.1 | $n_{S,m_s}$ | 1 | $S^0_{m_s}$ | 200 |
| **Activation on SNAIL by external signal I** | $\lambda_{I,m_s}$ | 10 | $n_{I,m_s}$ | 2 | $I^0_{m_s}$ | 50 |
| **Inhibition on GRHL2 by ZEB** | $\lambda_{Z,m_G}$ | 0.5 | $n_{Z,m_G}$ | 3 | $Z^0_{m_G}$ | 10 |
| **Self-activation of GRHL2** | $\lambda_{G,m_G}$ | 2 | $n_{G,m_G}$ | 2 | $G^0_{m_G}$ | 40 |

**Table SI 2. List of parameters for function $Y$ and $L$.**

| **n (# of miRNA binding sites)** | **0** | **1** | **2** | **3** | **4** | **5** | **6** |
|---|---|---|---|---|---|---|---|
| $l_i$**(hour$^{-1}$)** | 1 | 0.6 | 0.3 | 0.1 | 0.05 | 0.05 | 0.05 |
| $\gamma_{mi}$**(hour$^{-1}$)** | 0 | 0.04 | 0.2 | 1 | 1 | 1 | 1 |
| $\gamma_{\mu i}$**(hour$^{-1}$)** | 0 | 0.005 | 0.05 | 0.5 | 0.5 | 0.5 | 0.5 |
| $n_{\mu_{200}}$ | 6 | | | $n_{\mu_{34}}$ | | | 2 |
| $\mu^0_{200}$ | 10K | | | $\mu^0_{34}$ | | | 10K |

**Table SI 3. List of other parameters used in EMT model.**

| **Synthesis rate** | **Value (molecules/hour)** | **Degradation rate** | **Value (hour$^{-1}$)** | **Translation rate** | **Value (hour$^{-1}$)** |
|---|---|---|---|---|---|
| $g_{\mu_{200}}$ | 2.1K | $k_{\mu_{200}}$ | 0.05 | $g_z$ | 0.1K |
| $g_{m_z}$ | 11 | $k_{m_z}$ | 0.5 | $g_s$ | 0.1K |
| $g_{\mu_{34}}$ | 1.35K | $k_z$ | 0.1 | $g_G$ | 0.2K |
| $g_{m_s}$ | 90 | $k_{\mu_{34}}$ | 0.05 | | |
| $g_{mG}$ | 22 | $k_{m_s}$ | 0.5 | | |
| | | $k_s$ | 0.125 | | |
| | | $k_{mG}$ | 0.5 | | |
| | | $k_G$ | 0.1 | | |

### 5. More results about epigenetic feedback

#### Epigenetic feedback on GRHL2's self-activation does not greatly affect the EMT in reduced circuit

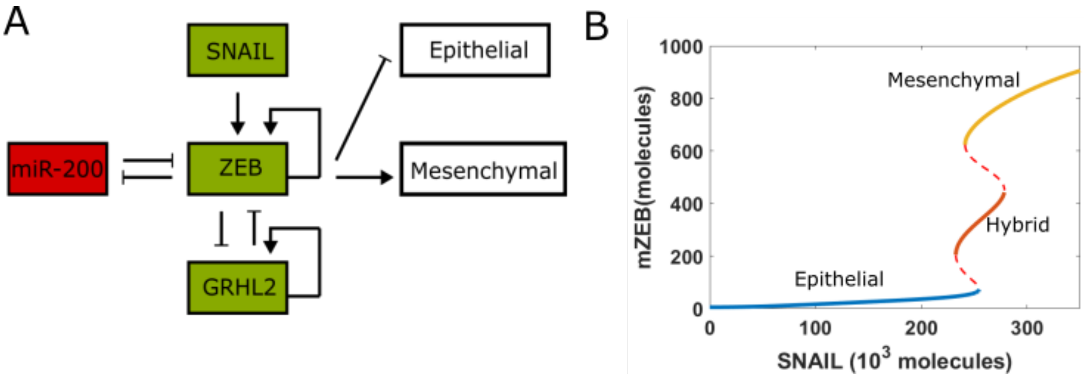


***Figure S4: Reduced EMT network*** *(A) The reduced core gene network regulates EMT – miR-200/ZEB mutually inhibitory circuit, coupled with GRHL2 and driven by SNAIL that serves as EMT-inducing signal here. (B) Bifurcation diagram of ZEB mRNA levels for the reduced network with SNAIL as the bifurcation parameter. Solid lines represent stable states, i.e. epithelial, hybrid or mesenchymal state, and dashed lines represent unstable states.*

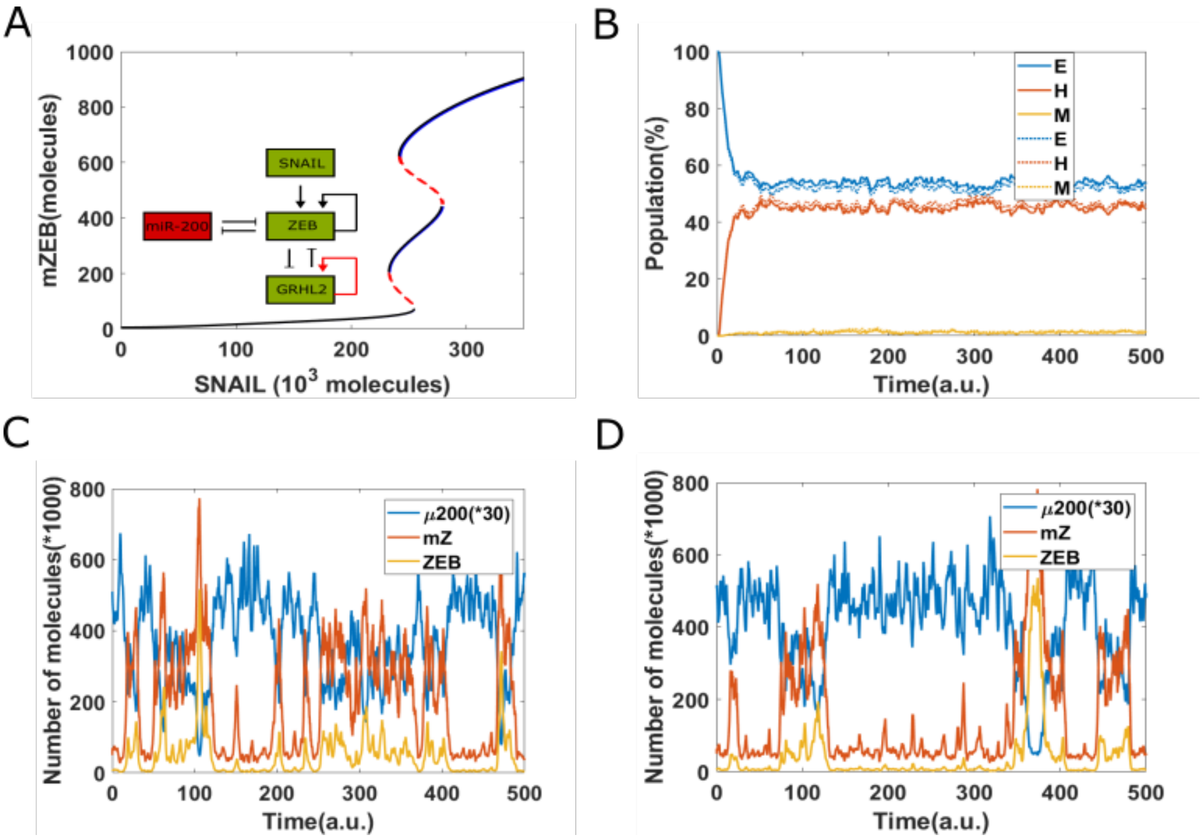

***Figure S5: Epigenetic feedback on GRHL2 self-activation for the reduced EMT network.*** *(A) The bifurcation diagrams for core EMT circuit with/without epigenetic feedback on self -activation of GRHL2. Black curves represent the non-epigenetic case. Blue curves represent the epigenetic case, and overlaps with the non-epigenetic curves. (B) Starting from pure epithelial state (miR-200=17,000, mZEB=50, ZEB=10,000 molecules), simulation results showing how the population changes as a function of simulation time. Dashed lines represent no epigenetic feedback case, and solid lines represent case with strong epigenetic feedback ($\alpha = 0.22$) on GRHL2's self-activation (SNAIL=245,000 molecules). (C) A sample dynamical diagram for no feedback case. (D) A sample dynamical diagram for strong feedback case.*

Our previous results suggested that the two mutually inhibitory feedback loops – miR-200/ZEB and miR-34/SNAIL – had two different roles in EMT decision-making. While miR-34/SNAIL loop was mostly monostable and acted as a noise-buffering integrator of various EMT/MET inducers, the miR-200/ZEB loop exhibited tristability and acted as a decision-making switch among the cells attaining one of the three possible phenotypes – epithelial (high miR-200, low ZEB), mesenchymal (low miR-200, high ZEB) and hybrid epithelial/mesenchymal (medium miR-200, medium ZEB) (3). Thus, we also investigated the dynamics of miR-200/ZEB circuit driven by SNAIL as an external signal (Fig S4A). The bifurcation diagram (Fig S4B) shows striking similarity to that of the network containing miR-34 (Fig 3B) - three distinct stable states are observed, i.e. epithelial(E), hybrid(H) and mesenchymal(M), which can co-exist within certain range of values of SNAIL levels.

Based on the results for the circuit including miR-34 (Fig 4), we quantified the effect of epigenetic feedback on self-activation of GRHL2 (Fig S5A). Similar results were obtained here as well – even when the epigenetic feedback is strong, the bifurcation diagram rarely changes (overlapping blue and black curves in Fig S5A). The population distribution obtained for SNAIL =245 x $10^3$ molecules, with all cells initially in an epithelial phenotype, was obtained to be approximately 54% E, 44% H and 2% M either with or without the epigenetic feedback (Fig S5B). Representative stochastic trajectories show that under the influence of noise, spontaneous transitions were seen among all the three phenotypes in both scenarios – with/without any epigenetic feedback (Fig S5C, D).

**Epigenetic feedback on the inhibition of GRHL2 by ZEB can stabilize epithelial state**

Next, for the reduced EMT network, we investigated the impact of epigenetic feedback on the inhibition of ZEB by GRHL2. Compared to the bifurcation diagram for no epigenetic feedback case (black curves), that of the strong epigenetic feedback case (blue lines) has shifted towards right, i.e. a higher external signal level is required to induce EMT; in other words, epithelial state is relatively more stabilized in presence of this epigenetic feedback (Fig S6A). Consistently, the population distribution is titled towards a higher relative percentage of epithelial cells; in presence of the epigenetic feedback, the stable population distribution is 80% E, 20% H and 0% M. Thus, compared to the no feedback case for the reduced network, the presence of this epigenetic feedback increases the percentage of epithelial cells by 26%, at the cost of reduced percentages for H and M states (Fig S6B). The representative stochastic trajectories also show that when there is strong epigenetic feedback on ZEB's inhibition from GRHL2, it is extremely rare for a cell to reach the mesenchymal state, and thus it usually stays much longer in the epithelial state (Fig S6C,D). Thus, these results show that the epigenetic feedback on GRHL2's self-activation does not affect the EMT/MET dynamics significantly, but the epigenetic feedback on inhibition of ZEB by GRHL2 can stabilize an epithelial state, irrespective of the presence or absence of miR-34/ SNAIL loop.

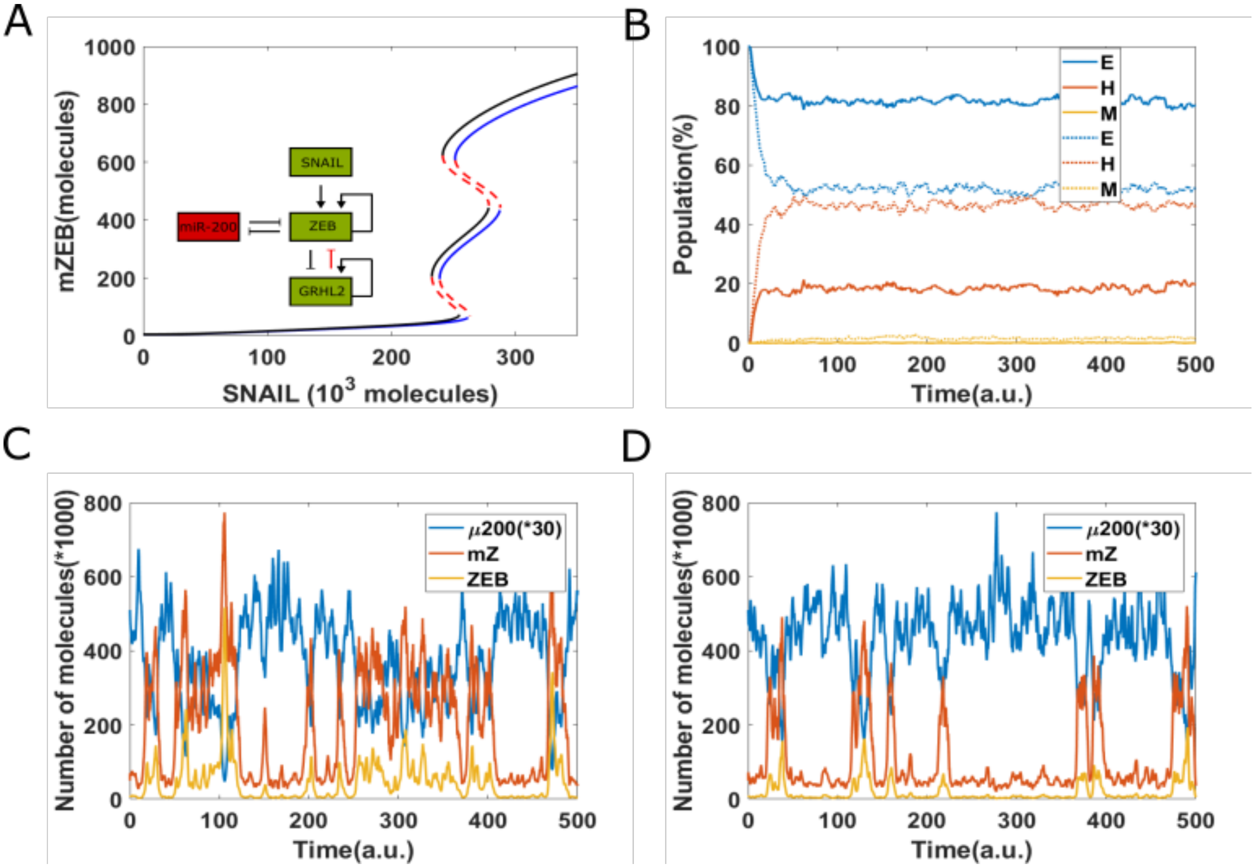


***Figure S6: Epigenetic feedback on inhibition of ZEB by GRHL2 in case of reduced network.*** *(A) The bifurcation diagrams for reduced EMT circuit with/without epigenetic feedback on the inhibition of ZEB by GRHL2. Black curves represent the non-epigenetic case. Blue curves represent the epigenetic case. (B) Starting from pure epithelial state (miR-200=17,000, mZEB=50, ZEB=10,000 molecules), simulation results showing how the population changes as a function of simulation time. Dashed lines represent no epigenetic feedback case, and solid lines represent case with strong epigenetic feedback* ($\alpha = 0.14$) *on ZEB's inhibition from GRHL2 (SNAIL=245,000 molecules). (C) A sample dynamical diagram for no feedback case. (D) A sample dynamical diagram for strong feedback case.*

**A Population-level Model of Epithelial-Mesenchymal Plasticity**

For the analysis shown in Fig 7, we used the model described previously by Tripathi *et al.* (7). We considered a population of tumor cells where each cell is carrying a copy of the core circuit regulating EMT / MET (shown in Fig 3 A). The dynamics of this circuit can be simulated for each cell using ordinary differential equations, independent of the other cells in the population.

Depending on the level of ZEB mRNA in a cell, it was assigned one of the three phenotypic states— epithelial (E), hybrid E/M, or mesenchymal (M). The stochastic population level dynamics was simulated using Gillespie's algorithm (8) and included two types of events: cell division and cell death. When a cell divides, the biomolecules (RNAs and proteins) are stochastically partitioned among the daughter cells. We only considered the stochastic partitioning of the EMT inducer $I_{sig}$ since it is likely to be the dominant perturbation to EMT / MET dynamics. Further, to incorporate the stem-like behavior of hybrid E/M cells such as asymmetric partitioning of miR-34a during cell division (9,10,11), whenever a hybrid E / M cells divided, one daughter cell got the same miR-34 concentration as the parent cell while concentration of miR-34 in the other daughter cell was set to 0. See Tripathi *et al.*(7) for further details regarding the population-level model.

The simulation whose results are shown in Fig 7 includes the following steps:

1. Start with a population of 500 cells where the level of the EMT inducer in each cells is drawn from a log normal distribution with median $2 \times 10^4$ molecules / cells and coefficient of variation 1.0. The dynamics of this population is simulated for a period of 28 days with a fixed carrying capacity of 10000. This step simulates the dynamics *in vivo.*
2. Carry out *in silico* FACS to obtain a population of 100 mesenchymal cells.
3. Simulate the dynamics of the population of mesenchymal cells obtained in step 2 with a carrying capacity of 500.
4. For the first 10 days (days 1-10), at the end of each day, a fixed dosage ($1 \times 10^4$ molecules / cell) of the EMT inducer ($I_{sig}$) was withdrawn from each cell in the population. If the concentration of $I_{sig}$ in a cell became negative during this process, it was set to 0. This step corresponds to cells in the population being treated so as to undergo MET.
5. For the next 10 days (days 11-20), at the end of each day, a fixed dosage ($1 \times 10^4$ molecules / cell) of the EMT inducer ($I_{sig}$) was added to each cell in the population. This step corresponds to cells in the population being treated so as to undergo EMT.

This simulation was carried out with and without GRHL2 activity. In both cases, 16 independent simulation runs were carried out and the mean and standard deviation over these runs has been shown in Fig 7.

### 6. References

[1] Cieply, B., Riley, P., Pifer, P. M., Widmeyer, J., Addison, J. B., Ivanov, A. V., ... & Frisch, S. M. (2012). Suppression of the epithelial–mesenchymal transition by Grainyhead-like-2. *Cancer research*, *72*(9), 2440-2453.

[2] Cieply, B., Farris, J., Denvir, J., Ford, H. L., & Frisch, S. M. (2013). Epithelial–mesenchymal transition and tumor suppression are controlled by a reciprocal feedback loop between ZEB1 and Grainyhead-like-2. *Cancer research*, *73*(20), 6299-6309.

[3] Lu, M., Jolly, M. K., Levine, H., Onuchic, J. N., & Ben-Jacob, E. (2013). MicroRNA-based regulation of epithelial–hybrid–mesenchymal fate determination. *Proceedings of the National Academy of Sciences*, *110*(45), 18144-18149.

[4] Jolly, M. K., Tripathi, S. C., Jia, D., Mooney, S. M., Celiktas, M., Hanash, S. M., ... & Levine, H. (2016). Stability of the hybrid epithelial/mesenchymal phenotype. *Oncotarget*, *7*(19), 27067.
[5] Levine, E., Jacob, E. B., & Levine, H. (2007). Target-specific and global effectors in gene regulation by MicroRNA. *Biophysical journal*, *93*(11), L52-L54.
[6] Lu, M., Jolly, M. K., Gomoto, R., Huang, B., Onuchic, J., & Ben-Jacob, E. (2013). Tristability in cancer-associated microRNA-TF chimera toggle switch. *The journal of physical chemistry B*, *117*(42), 13164-13174.
[7] Tripathi, S., Chakraborty, P., Levine, H., & Jolly, M. K. (2020). A mechanism for epithelial-mesenchymal heterogeneity in a population of cancer cells. *PLoS computational biology*, *16*(2), e1007619.
[8] Gillespie, D. T. (1977). Exact stochastic simulation of coupled chemical reactions. *The journal of physical chemistry*, *81*(25), 2340-2361.
[9] Bu, P., Chen, K. Y., Chen, J. H., Wang, L., Walters, J., Shin, Y. J., ... & Li, J. (2013). A microRNA miR-34a-regulated bimodal switch targets Notch in colon cancer stem cells. *Cell stem cell*, *12*(5), 602-615.
[10] Bu, P., Wang, L., Chen, K. Y., Srinivasan, T., Murthy, P. K. L., Tung, K. L., ... & Lipkin, S. M. (2016). A miR-34a-Numb feedforward loop triggered by inflammation regulates asymmetric stem cell division in intestine and colon cancer. *Cell stem cell*, *18*(2), 189-202.
[11] Wang, L., Bu, P., Ai, Y., Srinivasan, T., Chen, H. J., Xiang, K., ... & Shen, X. (2016). A long non-coding RNA targets microRNA miR-34a to regulate colon cancer stem cell asymmetric division. *Elife*, *5*, e14620.
